# BMP7 promotes cardiomyocyte regeneration

**DOI:** 10.1101/2023.08.17.553475

**Authors:** Chiara Bongiovanni, Hanna Bueno-Levy, Denise Posadas Pena, Irene Del Bono, Simone Redaelli, Max Bergen, Silvia Da Pra, Francesca Sacchi, Carmen Miano, Stefano Boriati, Francesca Pontis, Donatella Romaniello, Martina Mazzeschi, Ilaria Petraroia, Riccardo Tassinari, Laura Kellerer, Mattia Lauriola, Carlo Ventura, Stephan Heermann, Gilbert Weidinger, Eldad Tzahor, Gabriele D’Uva

## Abstract

Zebrafish has a remarkable and lifelong ability for cardiac regeneration after severe damage, whereas mammals lose their innate capacity for heart regeneration during early postnatal development. This study aimed to investigate whether the decreased production of growth factors during postnatal mammalian development contributes to the exit of cardiomyocytes from the cell cycle and the reduction in cardiac regenerative ability.

We identified growth factors with declining expression levels during early postnatal life in the mouse model and assessed the pro-proliferative ability of these factors on neonatal murine primary cardiomyocytes *in vitro*. Our findings confirmed the previously reported pro-proliferative effects of NRG1, IL1b, RANKL, IGF2 and IL6, while also identifying novel potential pro-regenerative growth factors. Among them, BMP7 exhibited the most pronounced efficacy.

Bmp7 knockdown interfered with the proliferation of neonatal mouse cardiomyocytes in culture and adult bmp7 mutant zebrafish displayed reduced cardiomyocyte proliferation during heart regeneration, indicating that Bmp7 is crucial for cardiomyocyte proliferation in the regenerative stages of mouse and zebrafish hearts. Conversely, *bmp7* overexpression was sufficient to boost cardiomyocyte cycling in regenerating zebrafish hearts, while BMP7 administration stimulated mouse cardiomyocyte cycling at postnatal-day-7, when cardiomyocytes ceased to proliferate, and enhanced cardiomyocyte regeneration *in vivo* in adult mice following myocardial infarction.

Mechanistically, BMP7-induced proliferation was mediated by type I BMP receptors BMPR1A and ACVR1, and type II receptors ACVR2A and BMPR2. Downstream signalling involved SMAD5, ERK and AKT.

In conclusion, the administration of BMP7 holds promise as a strategy to stimulate heart regeneration following cardiac injury.

## Introduction

In mammals, heart injuries such as those induced by myocardial infarction (MI), result in substantial loss of cardiac muscle cells (cardiomyocytes), which are replaced by fibrotic scar tissue. This condition, coupled with the very limited regenerative capacity of the adult mammalian heart, often leads to heart failure [reviewed in (Benjamin et al., 2019; van Berlo and Molkentin, 2014; Bongiovanni et al., 2021; Eschenhagen et al., 2017; Sadek and Olson, 2020; Tzahor and Poss, 2017)]. Currently, effective treatments for cardiac injuries are lacking, highlighting the urgent need to develop therapeutic strategies for cardiac regeneration.

While a remarkable and lifelong capacity for cardiac regeneration has been documented in certain non-mammalian vertebrates, including zebrafish and some amphibians (Poss et al., 2002), a significant ability to regenerate the heart in several mammalian species has been documented only during embryonic/foetal development and shortly after birth (Bryant et al., 2015; Drenckhahn et al., 2008; Haubner et al., 2012; Porrello et al., 2011; Sampaio-Pinto et al., 2018; Ye et al., 2018; Zhu et al., 2018). Indeed, mammalian cardiac regenerative ability declines rapidly in early postnatal life. In the mouse model, by one week after birth, most cardiomyocytes have exited the cell cycle (Li et al., 1996; Soonpaa and Field, 1998), making the heart unable to regenerate following injuries and resulting in the formation of permanent scars (Porrello et al., 2011). In adult mammals, cardiomyocyte turnover is extremely low (Bergmann et al., 2009; Senyo et al., 2013), and insufficient to initiate cardiac regeneration.

Studies have shown that reversible dedifferentiation and proliferation of cardiomyocytes play a crucial role in promoting cardiac regeneration in adult zebrafish (Jopling et al., 2010; Kikuchi et al., 2010) and neonatal mice (Porrello et al., 2011). Furthermore, endogenous cardiomyocytes represent the primary source of the low rate of cardiomyocyte turnover in adult mammals (Senyo et al., 2013). Various micro- environmental factors, including neuregulin 1 (NRG1) (Bersell et al., 2009; D’Uva et al., 2015a), fibroblast growth factor 1 (FGF1) (Engel et al., 2006), bone morphogenetic protein 10 (BMP10) (Sun et al., 2014), oncostatin M (OSM) (Kubin et al., 2011; Li et al., 2020a), tweak (Novoyatleva et al., 2010), T-reg secreted cytokines (Zacchigna et al., 2018a), follistatin (Wei et al., 2015), vascular endothelial growth factor (VEGF) (Tao et al., 2011), bone morphogenetic protein 1.3 (BMP1.3) (Vukicevic et al., 2022), bone morphogenetic protein 2 (BMP2) (Chakraborty et al., 2013; Ebelt et al., 2013), as well as several systemic factors like thyroid hormones (Hirose et al., 2019) and glucocorticoids (Pianca et al., 2022), have been shown to modulate dedifferentiation and proliferation of endogenous cardiomyocytes, thereby modulating heart regeneration in adult mammals following severe cardiac injuries (reviewed in (Bongiovanni et al., 2021; Cahill et al., 2017; Galdos et al., 2017; Hashimoto et al., 2018; Heallen et al., 2019; Sadek and Olson, 2020; Tzahor and Poss, 2017; Uygur and Lee, 2016)). Here, we hypothesise that the decline in expression levels of specific growth factors contributes to the loss of cardiomyocyte proliferation and regenerative ability in mammals soon after birth.

## Results

### The expression levels of several growth factors rapidly decline in heart tissue during early postnatal mammalian development

By employing the mouse model, we identified growth factors that exhibited the most pronounced decline in expression levels in heart tissue during early postnatal development, coinciding with the exit of cardiomyocytes from the cell cycle. To accomplish this, we examined a panel of 117 commercially available growth factors (**Supplementary Table 1**) in RNA-sequencing data obtained from hearts isolated from 1-day-old (P1) and 10-days-old (P10) mice (Haubner et al., 2012). Our analysis revealed that several growth factors display a decrease in expression levels during early postnatal life (**Fig. 1a**). Notably, 21 growth factors showed a reduction of over 50%, with 12 of them exhibiting a statistically significant decrease.

**Figure 1.**
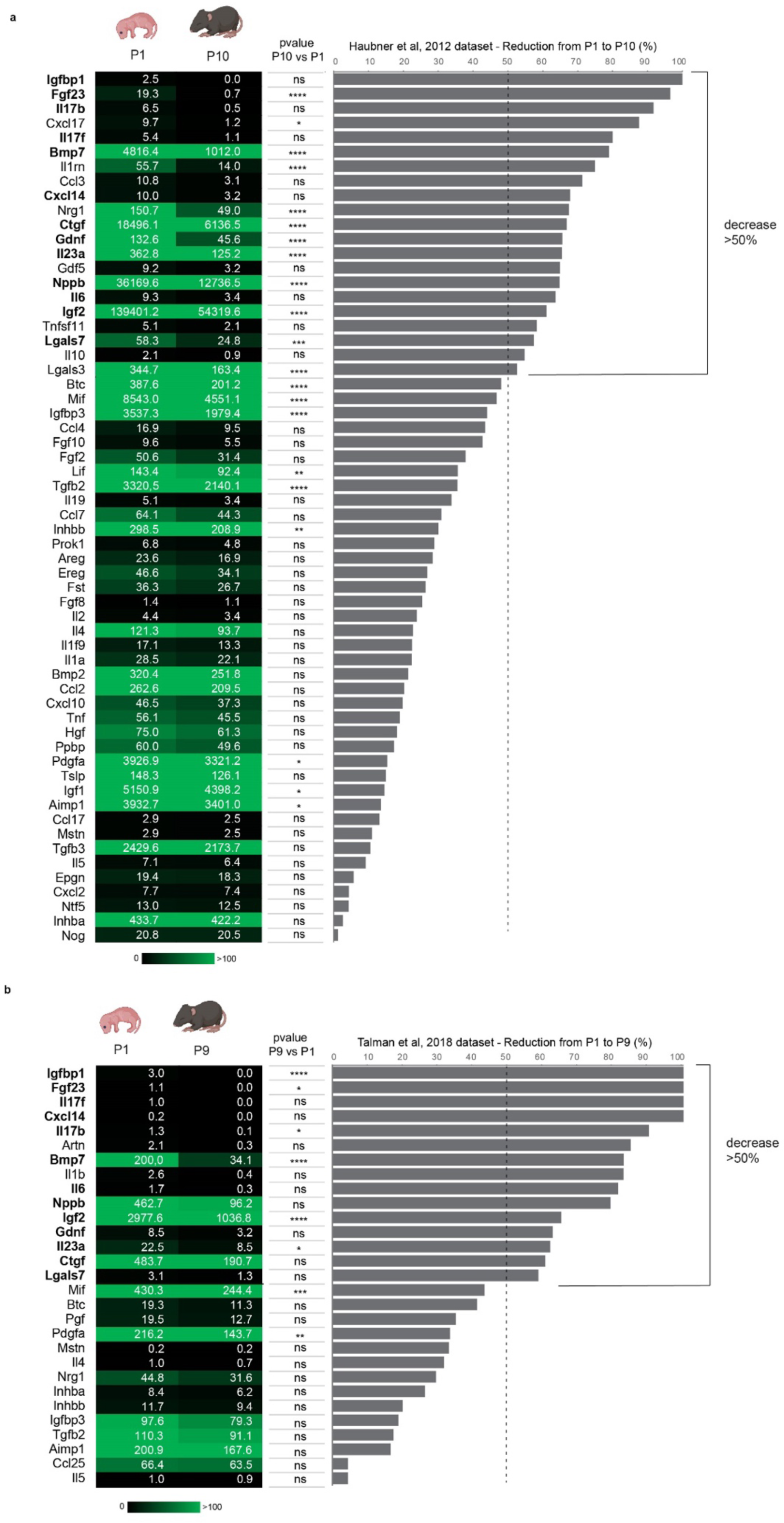
Identification of growth factors with declining expression levels during early cardiac postnatal development. (**a-b**) mRNA expression levels of growth factors in mouse postnatal day 1 (P1, first column) and postnatal day 9 or 10 (P9 or P10, second column) heart lysates obtained by meta-analysis of RNA-sequencing data ((Haubner et al., 2012) in **a**; (Talman et al., 2018) in **b**), along with the calculated decrease from P1 to P10 (or P9) in terms of percentage (graph bar). The values in the first and second columns are presented as mean expression levels of 3 biological replicates; the p-value is reported for the reduction from P1 to P10 (or P9) (third column) as follows * p ≤ 0.05; ** p ≤ 0.01; *** p ≤ 0.001; **** p ≤ 0.0001.

We conducted a similar analysis on a second dataset comprising RNA-sequencing data from hearts isolated from 1-day-old (P1) and 9-days-old (P9) mice (Talman et al., 2018). This analysis revealed a reduction of at least 50% in expression levels of 15 growth factors (**Fig. 1b**), 13 of which were also identified in the first analysis. By merging the results of the two datasets, we identified a total of 23 factors. Among these, genes encoding BMP7 (bone morphogenetic protein 7), IL1RA (IL1RN, interleukin 1 receptor antagonist), NRG1b (neuregulin 1b), CTGF (connective tissue growth factor), GDNF (glial cell line-derived neurotrophic factor), IL23a (interleukin 23a), BNP (NPPB, brain natriuretic peptide), IGF2 (insulin-like growth factor 2), LGALS7 (GAL7, galectin 7), LGALS3 (GAL3, galectin 3) were abundantly expressed at P1, while genes encoding for IGFBP1 (insulin-like growth factor binding protein), FGF23 (fibroblast growth factor 23), IL1b (interleukin 1b), IL17b (interleukin 17b), CXCL17 (C-X-C motif chemokine ligand 17), IL17f (interleukin 17f), CCL3 (C-C motif chemokine ligand 3), CXCL14 (C-X-C motif chemokine ligand 14), GDF5 (BMP14, bone morphogenetic protein 14), IL6 (interleukin 6), sRANKL (TNFSF1L, soluble RANK ligand) and IL10 (interleukin 10) had low expression levels at P1 (**Fig. 1a-b**).

### Several growth factors that exhibit a decline in expression during the early mammalian postnatal development promote cell cycle progression of neonatal cardiomyocytes

Subsequently, we evaluated the mitogenic properties of the 23 growth factors that demonstrated the most pronounced decline in expression during early postnatal development. We examined their effects on cell cycle progression of neonatal cardiomyocytes, which were isolated at postnatal day 1 (P1) and at this stage retain an intrinsic proliferative and regenerative ability reminiscent of the embryonic stage. Considering that the length of the cardiomyocyte S/G2/M phase during development has been estimated to be around 14 hours, with a total cell cycle length of over 24 hours (Hashimoto et al., 2014), we analysed the cell cycle progression of cardiomyocytes over a 48-hours timeframe using a cumulative BrdU incorporation assay, as previously done (Pianca et al., 2022). To specifically identify cardiomyocytes, we performed a co-immunostaining for the marker Troponin I (**Fig. 2**). Our data revealed that 18 out of 23 selected growth factors, namely CXCL17, sRANKL, BMP7, GDF5 (BMP14), LGALS7 (GAL7), CCL3, IGF2, IL6, IL1RA, IL17b, CXCL14, IL10, IGFBP, IL1b,

**Figure 2.**
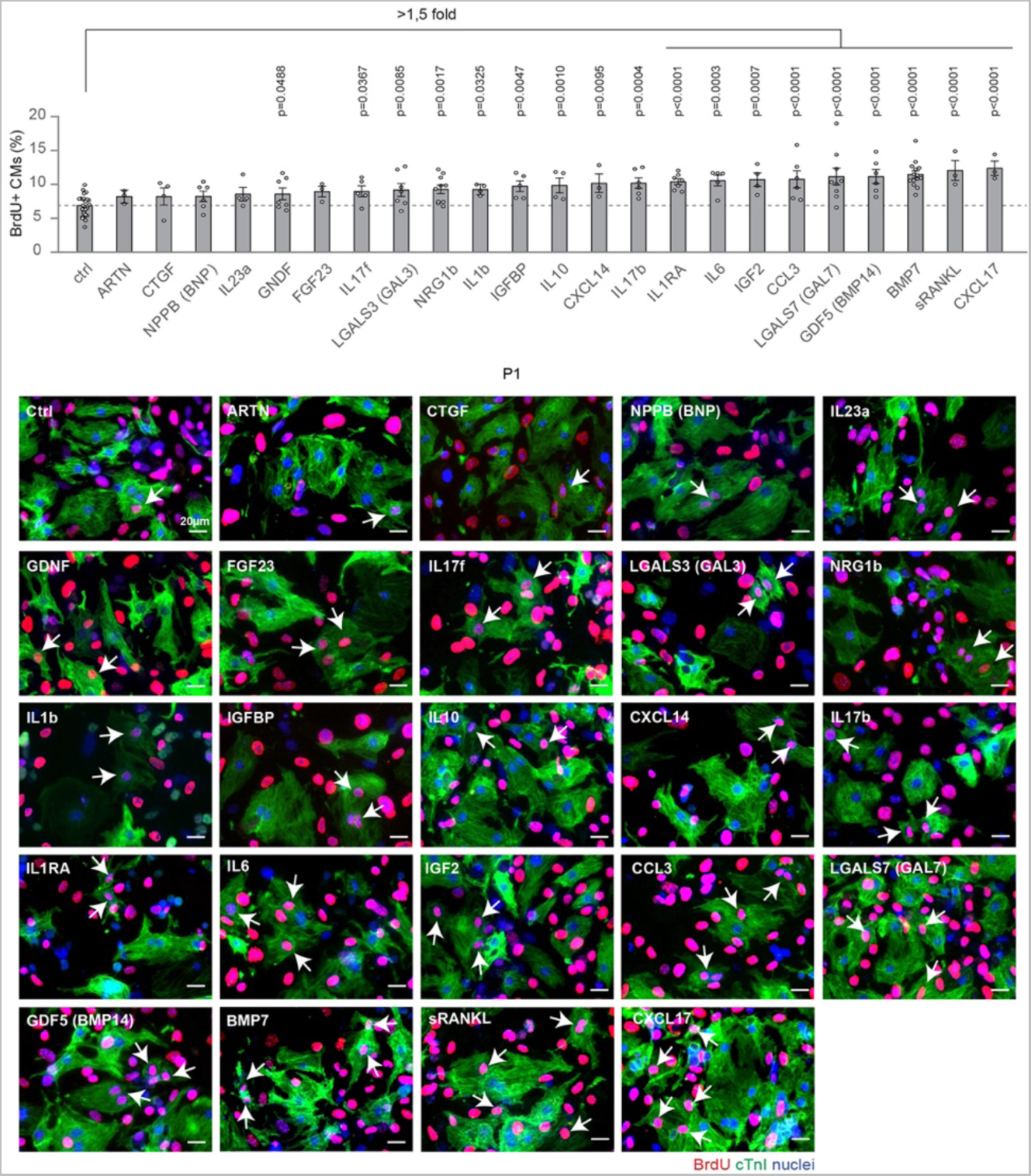
Ability of candidate growth factors to induce neonatal cardiomyocyte cell cycle progression. Cardiomyocytes isolated from neonatal mice were cultured in vitro and stimulated for 48 hours with selected growth factors (at the concentration of 10 ng/ml), namely ARTN, CTGF, NPPB (BNP), IL23a, GDNF, FGF23, IL17f, LGALS3 (GAL3), NRG1b, IL1b, IGFBP, IL10, CXCL14, IL17b, IL1RA, IL6, IGF2, CCL3, GDF5 (BMP14), LGALS7 (GAL7), BMP7, sRANKL, CXCL17. Cardiomyocytes were identified by cardiac Troponin I (cTnI) staining and analysed by immunofluorescence for DNA synthesis (BrdU incorporation assay) (n = 34914 cardiomyocytes pooled from the analysis of 162 samples); representative pictures are provided; arrows point at cardiomyocytes progressing in the cell cycle; scale bars, 20 µm. The values are presented as mean (error bars show s.e.m.), statistical significance was determined using one way ANOVA followed by Sidak’s test by comparing pairs of treatments (control vs selected growth factor).

NRG1b, LGALS3 (GAL3), IL17f and GDNF, were able to stimulate the progression of cardiomyocyte cell cycle into the S-phase (**Fig. 2**). Interestingly, NRG1 (Bersell et al., 2009; D’Uva et al., 2015a; Polizzotti et al., 2015), IGF2 (Shen et al., 2020; Zacchigna et al., 2018b), IL6 (Tang et al., 2018) and sRANKL (Zacchigna et al., 2018b) have previously been reported to promote cardiomyocyte proliferation and heart regeneration, validating our screening approach. IL1b has also been reported to induce cardiomyocyte proliferation (Palmer et al., 1995), although it may also lead to apoptosis (Li et al., 2020b). Importantly, our data unveiled the pro-proliferative potential of 13 novel factors. The treatment with CXCL17, BMP7, GDF5 (BMP14), LGALS7 (GAL7), CCL3, and IL1RA produced the most pronounced effects, resulting in more than 1.5- fold increase in cardiomyocyte cell cycle progression compared to untreated controls. IL17b, CXCL14, IL10, IGFBP, LGALS3 (GAL3), IL17f and GDNF also significantly boosted cardiomyocyte cell cycle progression, albeit to a lesser extent than the previous group of factors (**Fig. 2**).

### BMP7 acts as an autocrine growth factor promoting mammalian cardiomyocyte proliferation at neonatal stage

Based on the screening and analyses described above, we selected six growth factors that exhibited remarkable effects in the BrdU assay: CXCL17, BMP7, LGALS7 (GAL7), GDF5 (BMP14), CCL3, and IL1RA. We did not further analyse IGF2, IL6, and sRANKL as their effectiveness in promoting cardiomyocyte proliferation and cardiac regeneration has already been established in previous studies (Tang et al., 2018; Zacchigna et al., 2018b).

As mentioned earlier, the majority of mammalian cardiomyocytes exits the cell cycle shortly after birth(Li et al., 1996; Soonpaa and Field, 1998). To evaluate the ability of the selected growth factors to promote cell cycle activity in cardiomyocytes, we analysed the nuclear immunoreactivity for KI67, a marker for the active phases of the cell cycle and used cardiac Troponin T as a marker for cardiomyocytes. Our results confirmed that all of the selected growth factors stimulate cardiomyocyte cell cycle activity, with BMP14 and BMP7, both belonging to the bone morphogenetic protein family, exhibiting the most robust effects (**Fig. 3a**).

**Figure 3.**
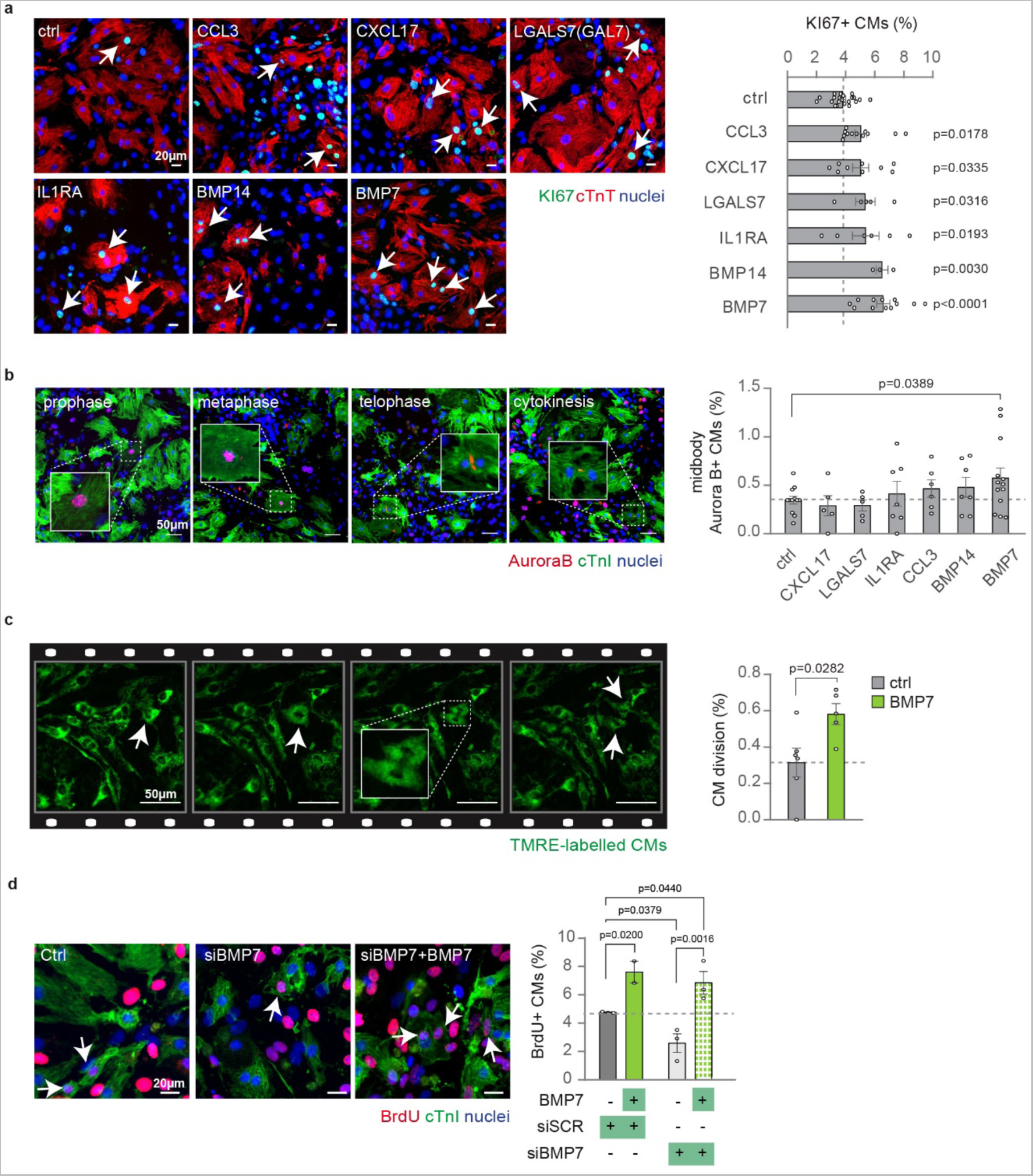
BMP7 robustly induces cell cycle activity and cell division in neonatal mammalian cardiomyocytes. **(a-b)** Cardiomyocytes isolated from post-natal day 1 (P1) mice were cultured *in vitro* and stimulated for 48 hours with selected growth factors (at the concentration of 10 ng/ml), namely CCL3, CXCL17, GAL7 (LSGAL7), IL1RA, BMP14 and BMP7. Cardiomyocytes were identified by cardiac Troponin T (cTnT) or Troponin I (cTnI) staining and analysed by immunofluorescence for (**a**) cell-cycle activity (KI67) or (**b**) cytokinesis (midbody Aurora B kinase) (n = 13961 cardiomyocytes pooled from the analysis of 68 samples in **a**; n = 57498 cardiomyocytes pooled from the analysis of 55 samples in **b**); representative pictures are provided; arrows point at cycling cardiomyocytes; scale bars, 20 µm in **a** and 50 µm in **b**; **(c)** Quantification and representative images of cell division events (n = 11 samples with a total of 5184 cardiomyocytes analysed) in TMRE-labelled neonatal cardiomyocytes (see Methods for details) detected in 16-hour time-lapse imaging at 15-minute intervals *in vitro*; arrows point at cardiomyocytes undergoing cell division; scale bars, 50 μm. **(d)** Evaluation of neonatal (postnatal day 1, P1) cardiomyocyte proliferation by immunofluorescence analysis of DNA synthesis (BrdU assay) following knockdown of BMP7 with and without BMP7 stimulation at 10 ng/ml for 48 hours (n = 2480 cardiomyocytes pooled from the analysis of 11 samples). Representative pictures are provided; scale bars 20 µm; arrows point at cycling cardiomyocytes. The values are presented as mean (error bars show s.e.m.), statistical significance was determined using one way ANOVA followed by Sidak’s test in **a**, **b** and **d** (comparison between pairs of treatments; control vs selected growth factor) and two-sided Student’s t-test in **c**.

During the early postnatal period in mice, most cardiomyocytes undergo DNA synthesis and nuclear division (karyokinesis) without proceeding to cytoplasm division (cytokinesis), resulting in binucleation (Soonpaa et al., 1996). Similarly, in humans, most cardiomyocytes undergo DNA synthesis without karyokinesis, resulting in polyploidization [reviewed in (Derks and Bergmann, 2020)]. To assess the impact of the selected growth factors on neonatal cardiomyocyte division, we analysed the staining for Aurora B kinase, which localizes at the equator of the central spindle during late anaphase and at the midbody during cytokinesis.

Our data demonstrated that BMP7 has a significant effect in triggering cardiomyocyte cytokinesis (**Fig. 3b**). This effect was further confirmed by time-lapse imaging of neonatal cardiomyocytes labelled with tetramethylrhodamine ethyl ester (TMRE), a fluorescent mitochondrial dye (Hattori et al., 2010) (**Fig. 3c**).

By performing negative immuno-magnetic selection for cardiac stromal cells, the ability of BMP7 in promoting cardiomyocyte proliferation was also confirmed in enriched-cardiomyocyte cultures (**Supplementary Fig. 1a, b**). These data suggest that the effect of BMP7 on neonatal cardiomyocyte proliferation is direct.

To evaluate if endogenous BMP7 sustains cardiomyocyte proliferation at the neonatal stage we knocked down BMP7. The efficiency of BMP7 silencing was confirmed 48 hours post transfection (**Supplementary Fig. 2**). Our data show that BMP7 knockdown reduces the proliferation of cultured postnatal day 1 (P1) cardiomyocytes, which was restored by exogenous administration of BMP7 (**Fig. 3d**).

Overall, these data indicate that endogenous BMP7 directly promotes cardiomyocyte cell cycle activity, progression to the S-phase and complete cell division.

### Bmp7 promotes cardiomyocyte proliferation during the spontaneous cardiac regeneration process in zebrafish

The BMP7 protein is encoded by a gene that is highly conserved across vertebrate species (Dong et al., 2022; Shawi and Serluca, 2008). Recent studies have demonstrated the involvement of BMP signalling in the highly efficient process of cardiac regeneration following injury in adult zebrafish. Our previous research has shown that BMP signalling is activated after cardiac injury in zebrafish, and its inhibition prevents cardiomyocyte regeneration (Wu et al., 2016). While an increase in the expression of BMP ligands (*bmp2b* and *bmp7a*) has been documented upon injury (Wu et al., 2016), the specific role of Bmp7 in cardiomyocyte regeneration in zebrafish has not been previously explored.

To investigate the role of Bmp7 in zebrafish cardiomyocyte regeneration, we analysed regenerating hearts of zebrafish homozygous for a loss-of-function mutation in the *bmp7a* gene (Schmid et al., 2000). By injecting wild-type *bmp7a* mRNA into fertilized eggs, we were able to rescue the early embryonic lethality of homozygous mutants (Dong et al., 2022) and allow them to reach adulthood (**Fig. 4a**). We then assessed EdU incorporation at 7 days post cryoinjury (dpi), when cardiomyocyte proliferation is at its peak, and found that cardiomyocyte proliferation within the wound border zone was reduced in *bmp7a*-deficient fish compared to wild-type siblings (**Fig. 4b**).

**Figure 4.**
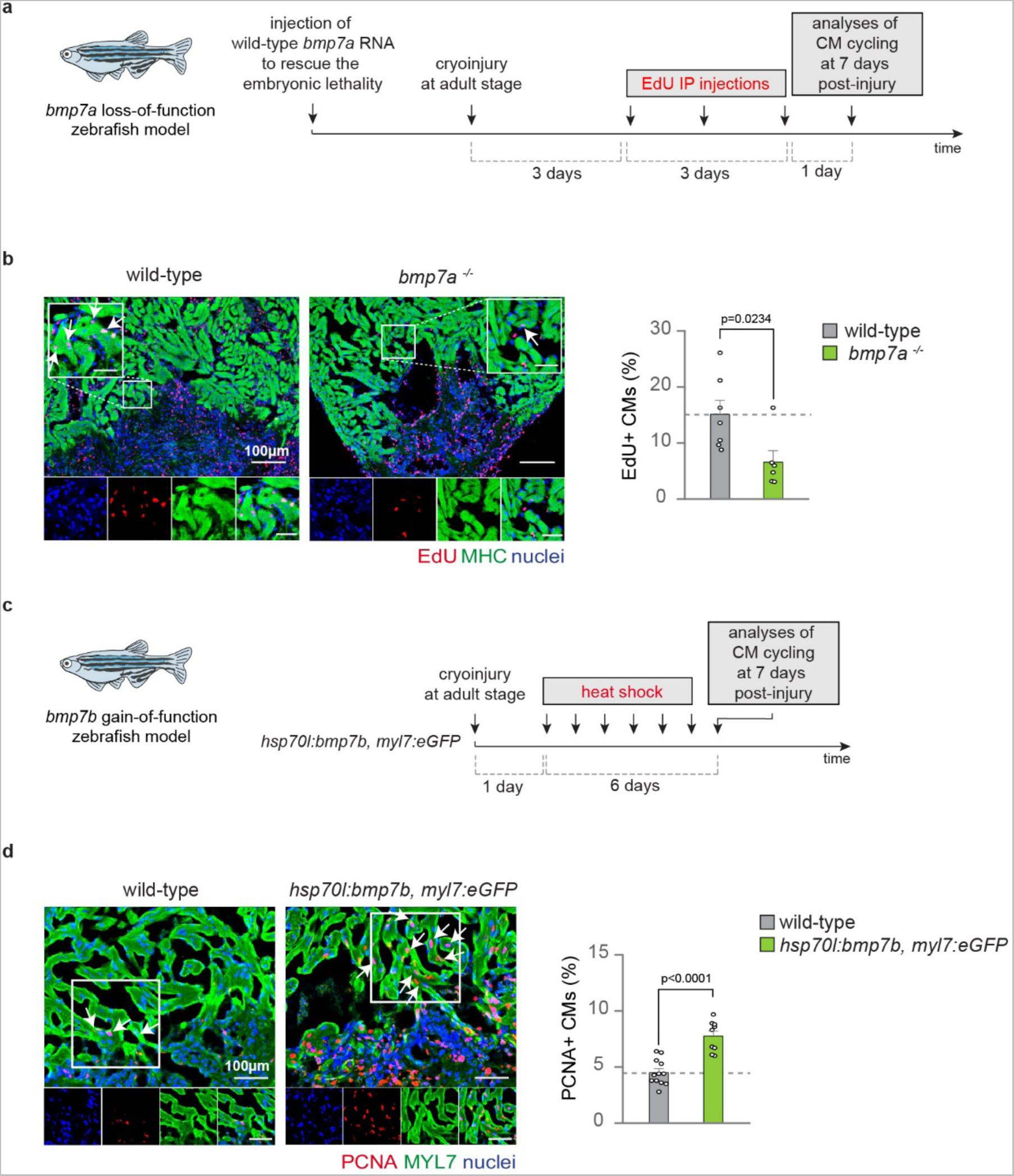
Bmp7 loss- and gain-of-function respectively reduces and increases cardiomyocyte proliferation during zebrafish heart regeneration. **(a)** Experimental design for the analysis of cardiomyocyte cell cycling at 7 days post cryoinjury (dpi) in *bmp7a* homozygous mutant fish; **(b)** EdU and MHC immunostaining (to identify cardiomyocytes) on cryoinjured hearts of *bmp7a* mutant fish and wild-type siblings. n (wild-type) = 7 hearts, 3 sections per heart, 1158 cardiomyocytes, n (*bmp7a^-/-^*) = 6 hearts, 3 sections per heart, 472 cardiomyocytes. Representative pictures are provided; arrows point at cycling cardiomyocytes; scale bars, 100 μm; **(c)** Experimental design depicting the use of a transgenic line allowing for heat-shock inducible overexpression of *bmp7b* together with constitutive expression of GFP in cardiomyocytes; (**d**) Immunofluorescence analysis of PCNA(proliferating cell nuclear antigen) and myosin light chain 7 (MYL7, to identify cardiomyocytes) in heat-shocked *hsp70l*:bmp7b, *myl7*:eGFP^af5Tg^ transgenic and wild type siblings at 7 dpi. Note that GFP fluorescence is not shown. n (wild-type) = 12 hearts, 2 sections per heart, 421 cardiomyocytes. n (*hsp70l*:bmp7b, *myl7*:eGFP) = 9 hearts, 2 sections per heart, 472 cardiomyocytes. Edu and PCNA positive cardiomyocytes in panel **b** and **d**, respectively, were counted manually within the border zone of control and transgenic fishes. Each dot represents a different heart (biological replicate), which is the average of the analysis of 2 to 3 sections. Representative pictures are provided; arrows point at cycling cardiomyocytes; scale bars, 100 μm. The values in **b, d** are presented as mean, error bars show s.e.m., statistical significance was determined using two-sided Student’s t-test.

The zebrafish genome contains two orthologs of human BMP7, *bmp7a* and *bmp7b*, with Bmp7b being more similar to human BMP7 than Bmp7a (∼80% compared to ∼66 % amino acid identity, respectively) (Shawi and Serluca, 2008). Previous studies have shown that the deletion of bmp ligands including *bmp2b*, *bmp4* and *bmp7* results in dorsalised phenotypes and early embryonic lethality (Dong et al., 2022), while their overexpression induces ventralisation (Schmid et al., 2000). We found that *bmp7a* and *bmp7b* were equally efficient in ventralising wild-type embryos (**Supplementary Fig. 3a-b**), and that *bmp7b* mRNA could fully rescue *bmp7a* mutant embryos (**Supplementary Fig. 3c-d; Supplementary Table 3**). We conclude that zebrafish *bmp7a* and *bmp7b* have similar biological activity in overexpression assays. We thus chose *bmp7b* for further gain-of-function experiments since its sequence is more similar to human BMP7. We generated a transgenic zebrafish line for heat-shock-inducible overexpression of *bmp7b* and concomitant labelling of cardiomyocytes by eGFP (*hsp70l*:bmp7b, *myl7*:eGFP^af5Tg^). In this transgenic line, *bmp7b* overexpression was induced by daily heat-shock starting at 1 dpi for 6 days (**Fig. 4c**), resulting in a significantly increased number of proliferating cardiomyocytes at 7 dpi compared to heat-shocked wild-type fish (**Fig. 4d**).

Together, these results demonstrate that endogenous *bmp7a* sustains cardiomyocyte proliferation during innate cardiac regeneration in zebrafish, and that further augmentation of *bmp7* levels can enhance this process (**Supplementary Fig. 4**).

### *Bmp7* expression levels are specifically reduced in cardiomyocytes during the postnatal cardiac mammalian development

We examined *Bmp7* expression in cardiac tissues obtained at different stages of postnatal murine development, namely P1, P3, P7, P28, and P56. Our analysis confirmed a significant decrease in *Bmp7* expression during the early postnatal development, and this decline continued beyond that stage (**Fig. 5a**). Using positive and negative immuno-magnetic selection for cardiac stromal cells, we also observed that *Bmp7* is more highly expressed in cardiomyocytes compared to cardiac stromal cells in neonatal (P1) mice, and that there is a specific reduction in expression within cardiomyocytes during early postnatal development (**Fig. 5b**). To better characterize *Bmp7* expression in the heart tissue, we analysed publicly available data on sorted cardiac cell populations (Quaife-Ryan et al., 2017). These analyses showed that *Bmp7* expression during the neonatal stage is primarily observed in cardiomyocytes and to a lesser extent in immune cells **(Supplementary Fig. 5a).** Conversely, neonatal fibroblasts and endothelial cells exhibited negligible levels **(Supplementary Fig. 5a).** Furthermore, *Bmp7* expression levels in cardiomyocytes dramatically declined from the neonatal to the adult stage and was not induced by neonatal or adult heart injury **(Supplementary Fig. 5b)**. Thus, BMP7 is predominantly produced by neonatal cardiomyocytes, and its expression decreases during early postnatal development.

**Figure 5.**
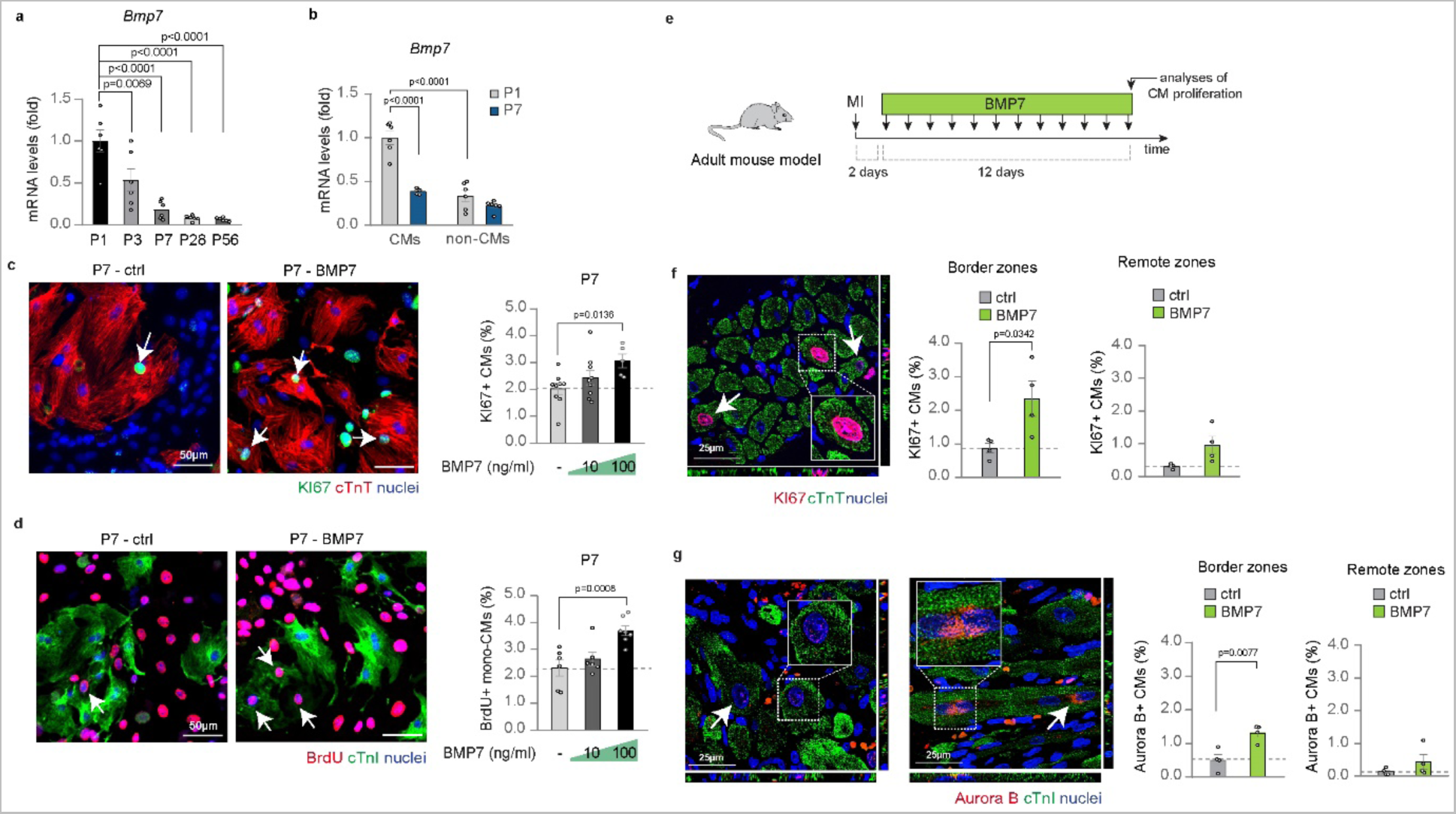
BMP7 administration stimulates the proliferation of mammalian cardiomyocytes at juvenile stage and in adult life following myocardial infarction. **(a)** *Bmp7* mRNA expression levels in heart lysates from postnatal day 1 (P1), 3 (P3), 7 (P7), 28 (P28) and postnatal day 56 (P56) mice (n = 6 hearts per developmental stage) as determined by qRT-PCR and shown relative to the level at P1; (b) *Bmp7* mRNA expression levels in cardiomyocytes and stromal cells isolated from postnatal day 1 (P1) and postnatal day 7 (P7) hearts and separated by immunomagnetic separation (n = 10 samples for cardiomyocytes and 12 for stromal cells); (c) *In vitro* evaluation of the impact of BMP7 on post-mitotic cardiomyocyte cell-cycle re-entry by immunofluorescence analysis of KI67 and cardiac Troponin T (cTnT) in postnatal day 7 (P7) cardiomyocytes following stimulation with BMP7 at 10 and 100 ng/mL for 48 hours (n = 5553 cardiomyocytes pooled from the analysis of 23 samples); Representative pictures are provided; arrows point at cycling cardiomyocytes; scale bars 50 µm; (d) *In vitro* evaluation of the impact of BMP7 on proliferation of postnatal-day-7 (P7) cardiomyocytes by immunofluorescence analysis of BrdU incorporation and cardiac Troponin I (cTnI) in postnatal day 7 (P7) mononucleated cardiomyocytes following stimulation with BMP7 at 10 and 100 ng/mL for 48 hours (n = 2843 cardiomyocytes pooled from the analysis of 19 samples). Representative pictures are provided; arrows point at proliferating cardiomyocytes; scale bars 50 µm; **(e)** Experimental design for the analysis of cardiac regeneration following ligation of the left anterior descending coronary artery in adult mice (3 months old); (f-g) *In vivo* evaluation of adult cardiomyocyte proliferation by immunofluorescence analysis of (**f**) KI67 and cardiac Troponin T (cTnT) or (**g**) Aurora B kinase and cardiac Troponin I (cTnI) in the border and remote zones of heart sections 14 days post myocardial infarction, following daily injection of BMP7 or water as control (n = 8 mice; for KI67/cTnT staining a total of 14345 cardiomyocytes have been analysed in the border zones and 23085 cardiomyocytes in the remote zones; for Aurora B/cTnI staining a total of 10088 cardiomyocytes have been analysed in the border zones and 14242 cardiomyocytes in the remote zones). KI67 and Aurora B kinase positive cardiomyocytes in tissue sections were counted manually within border or remote zones. Each dot represents a different heart (biological replicate), which is the average of the analysis of 2 to 3 sections. Representative pictures show images taken by confocal microscopy showing the xy maximum intensity projection of the stacks acquired from the sample (the slices view on the bottom and left of each panel represent the xz and yz stacks respectively); arrows point at proliferating cardiomyocytes; scale bars 25 µm. The values in **a**, **b, c, d**, **f** and **g** are presented as mean (error bars show s.e.m.); statistical significance was determined using one way ANOVA followed by Sidak’s test in **a, b, c** and **d** (comparison between pairs of treatments; control vs BMP7) or using two-sided Student’s t-test in **f-g**.

Overall, our data suggest that BMP7 may act as an autocrine factor in cardiomyocytes, triggering their proliferation.

### Administration of BMP7 stimulates cardiomyocyte proliferation after myocardial infarction in adult mammals

In mammals, the regenerative ability of cardiomyocytes is dramatically reduced during the early postnatal life (Porrello et al., 2011), coincident with their maturation, binucleation and cell cycle exit (Soonpaa et al., 1996). To investigate whether BMP7 signalling can induce proliferation in cardiomyocytes after the closure of the regenerative window, we examined the impact of BMP7 treatment at varying concentrations on cardiomyocytes isolated from postnatal day 7 (P7) mice. Our results show that high doses of BMP7 result in an increase in cell cycle activity (as indicated by KI67 staining) and cell cycle progression (as indicated by BrdU assay) in P7 cardiomyocytes (**Fig. 5c**; **Supplementary Fig. 6a**). Additionally, following high-dose BMP7 administration, there was a robust increase in the number of BrdU+ mononucleated cardiomyocytes (**Fig. 5d**) accompanied by a tendency towards a reduction in BrdU+ binucleated cardiomyocytes (**Supplementary Fig. 6b**). These observations suggest that BMP7 is sufficient to stimulate cell division also in cardiomyocytes at juvenile stage, when the regenerative ability is dramatically diminished.

Interestingly, a recent study has shown that BMP7 administration for two weeks after myocardial infarction (MI) has cardioprotective effects, reducing infarct size and improving cardiac function in rats (Jin et al., 2018). The suggested mechanism involves the counteraction of TGF-β1 pro-fibrotic signalling pathway (Jin et al., 2018). However, the potential of BMP7 to promote the regeneration of adult mammalian cardiomyocytes after a cardiac injury has not been evaluated.

To assess whether BMP7 can stimulate cardiomyocyte proliferation after cardiac injury, we induced myocardial infarction (MI) by permanently ligating the left anterior descending coronary artery in adult mice (3 months old), and we administered BMP7 for 12 days through intravenous/intraperitoneal injections, starting 2 days after MI (**Fig. 5e**). At the end of the treatment (14 days post-MI), we observed an increase in cell cycle re-entry and proliferation of cycling cardiomyocytes in infarct border zone (**Fig. 5f-g**, **Supplementary Fig. 6c**), with a trend towards an increase in remote zones (**Fig. 5f-g**). Importantly, our results suggest that the pro-proliferative effect of BMP7 is specific to cardiomyocytes, as there was a modest decrease in cycling cardiac stromal cells in BMP7-treated infarcted mice **(Supplementary Fig. 7a)**. The anti-proliferative effect of BMP7 on cardiac stromal cells was also observed in neonatal cardiac cell cultures **(Supplementary Fig. 7b-c)**. Overall, our data suggest that BMP7 is a growth factor capable of triggering mammalian cardiomyocyte proliferation. The decline in its abundance in the cardiac tissue during early postnatal mammalian development contributes to cardiomyocyte cell cycle exit, and its administration after myocardial infarction in adult mammals may represent a novel strategy for cardiac regeneration (**Supplementary Fig. 8**)

### BMP7 stimulates cardiomyocyte proliferation via BMPR1A/ACVR1 type I receptors and BMPR2/ACVR2A type II receptors

BMP7 is a member of the bone morphogenetic proteins (BMPs), which belong to the transforming growth factor-β (TGFβ) superfamily. The activities of all TGFβ family ligands are mediated by tetramers of serine/threonine kinase receptors, consisting of two type I and two type II receptors (Heldin and Moustakas, 2016; Loomans and Andl, 2016; Miyazono et al., 2005). BMPs have been reported to signal through ACVRL1, ACVR1, BMPR1A, ACVR1B, and BMPR1B as type I receptors, and BMPR2, ACVR2A and ACVR2B as type II receptors (Heldin and Moustakas, 2016; Loomans and Andl, 2016; Miyazono et al., 2005). Specifically, BMP7 has been shown to bind and activate ACVR1, BMPR1A, BMPR2, ACVR2A (Lavery et al., 2008), BMPR1B (González-Gómez et al., 2015), and ACVR2B (Yamashita et al., 1995) in different cell types and development stages.

To understand the molecular mechanism by which BMP7 regulates cardiomyocyte proliferation, we first analysed the expression levels of BMP receptors in neonatal murine cardiomyocytes using publicly available datasets (Quaife-Ryan et al., 2017). We found that *Bmpr1a* and *Bmpr2* were the most highly expressed type I and type II BMP receptors, respectively (**Supplementary Fig. 9a**). *Acvr1b*, *Acvrl1*, *Acvr1* and *Acvr2a* showed weak expression, whereas *Bmpr1b* and *Acvr2b* were not expressed (**Supplementary Fig. 9a**) and were excluded from further analyses.

To determine the receptors responsible for the proliferation induced by BMP7, we silenced the genes encoding BMPR1A, ACVR1, ACVRL1, ACVR1B, BMPR2 and ACVR2A in neonatal cultured cardiomyocytes, and then performed a BrdU incorporation assay following BMP7 stimulation. The efficiency of BMP receptor silencing was confirmed 48 hours post transfection (**Supplementary Fig. 9b**). The results showed that knockdown of BMPR1A, ACVR1, BMPR2 or ACVR2A abolishes the mitogenic effects of BMP7, while knockdown of ACVR1B and ACVRL1 had no significant effect (**Fig. 6a**). These findings suggest that BMP7 promotes cardiomyocyte proliferation through BMPR1A/ACVR1 and BMPR2/ACVR2A receptors, likely forming a tetramer as previously reported in human mesenchymal stem cells (Lavery et al., 2008).

**Figure 6.**
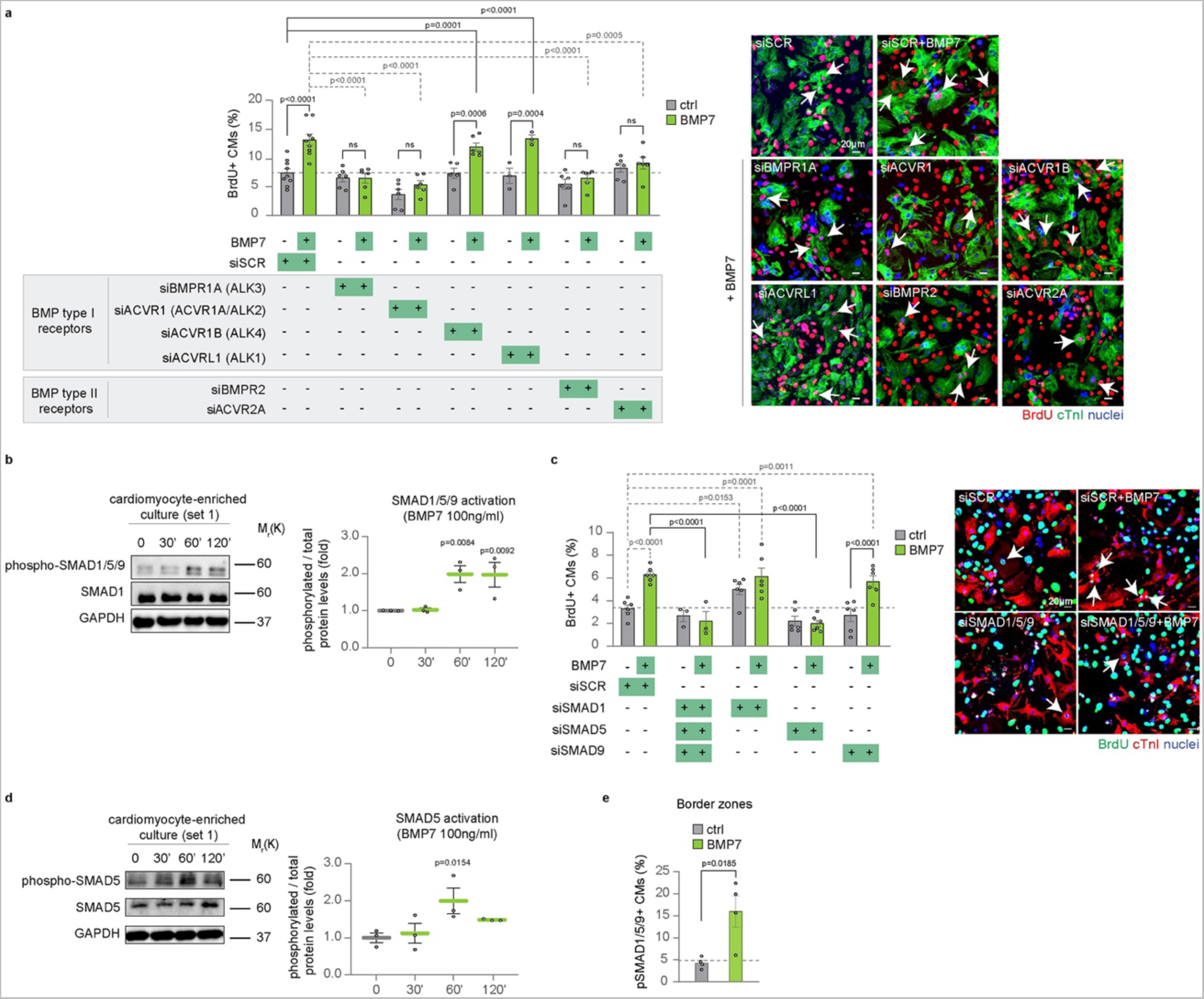
BMP7 induces cardiomyocyte proliferation through BMPR1A/ACVR1 and ACVR2/BMPR2 receptors and canonical SMAD5 signalling. **(a)** Evaluation of the role of BMP receptors in BMP7-induced cardiomyocyte proliferation by immunofluorescence analysis of DNA synthesis (BrdU assay) in neonatal (postnatal day 1, P1) cardiomyocytes following BMPR1A, ACVR1, ACVR1B, ACVRL1, BMPR2 and ACVR2A knockdown, upon BMP7 stimulation at 10 ng/ml for 48 hours (n = 23895 cardiomyocytes pooled from the analysis of 83 samples); representative pictures are provided; scale bars 20 µm, arrows point at proliferating cardiomyocytes; **(b)** Western Blot analysis of phospho-SMAD1/5/9, SMAD1 and GAPDH protein levels in enriched neonatal cardiomyocytes, separated from stromal cells by immunomagnetic separation, following BMP7 stimulation at 100 ng/ml for 30, 60 and 120 minutes (n = 3 replicates per condition). GAPDH protein levels are provided as second loading control; **(c)** Immunofluorescence analysis of DNA synthesis (BrdU assay) and cardiac Troponin I (cTnI) on neonatal (postnatal day 1, P1) cardiomyocytes following SMAD1, SMAD5 and SMAD9 knockdown, alone or in combination, upon BMP7 stimulation at 10 ng/ml for about 48 hours (n = 19167 cardiomyocytes pooled from the analysis of 54 samples); representative pictures are provided; scale bars 20 µm, arrows point at proliferating cardiomyocytes; **(d)** Western Blot analysis of phospho-SMAD5, SMAD5 and GAPDH protein levels in enriched neonatal (postnatal day 1, P1) cardiomyocytes, separated from stromal cells by immunomagnetic separation, following BMP7 stimulation at 100 ng/ml for 30, 60 and 120 minutes (n = 3 replicates per condition). GAPDH protein levels are provided as second loading control; (**e**) Immunofluorescence analysis for phospho-SMAD1/5/9 and cardiac Troponin T (cTnT) in the border zones of heart sections 14 days post myocardial infarction, following daily injection of BMP7 or water as control (n = 8 mice; a total of 8342 cardiomyocytes have been analysed). Phospho- SMAD1/5/9 positive cardiomyocytes in tissue sections were counted manually within border zones; every dot represents a different heart (biological replicate), which in turn has been calculated as average of the analysis of 1-2 sections. The values in **a-e** are presented as mean (error bars show s.e.m.); statistical significance was determined using one way ANOVA followed by Sidak’s test (comparison between pairs of treatments) in **a-d**; and using two-sided Student’s t-test in **e**.

### Canonical SMAD5 transduces the mitogenic signalling of BMP7

BMP ligands activate type II receptors, which phosphorylate the kinase domain of type I receptors and activate multiple downstream pathways (Heldin and Moustakas, 2016; Lavery et al., 2008). The canonical pathway involves receptor-regulated SMAD proteins (R-SMADs), including SMAD1, SMAD5, and SMAD8/9 for BMP-induced signalling. Once phosphorylated and activated, these SMAD proteins interact with the common factor SMAD4 and translocate into the nucleus (Massagué et al., 2005).

Therefore, we analysed the activation of canonical SMAD signalling in cultured neonatal cardiomyocytes after BMP7 stimulation. Our data showed a transient activation of SMAD1/5/9 in both cardiomyocyte-enriched cultures and cardiac cell cultures, as evidenced by higher levels of phosphorylated SMAD1/5/9 (p-SMAD1/5/9) compared to untreated controls (**Fig. 6b, Supplementary Fig. 10a-b**). The combined knockdown of SMAD1, SMAD5, and SMAD9 completely abolished the mitogenic effect of BMP7, indicating that SMAD1/5/9 signalling is necessary for BMP7-induced cardiomyocyte proliferation (**Fig. 6c**). Single or double R-SMADs can associate with SMAD4, thus resulting in heterodimeric or heterotrimeric complex (Massagué et al., 2005). We therefore investigated the effects of individual SMAD knockdown on cardiomyocyte proliferation after exposure to BMP7. The efficiency of SMAD silencing was confirmed 48 hours post transfection (**Supplementary Fig. 10c**). Unexpectedly, our data showed an opposite trend for SMAD1 and SMAD5 on cardiomyocyte proliferation. SMAD1 knockdown significantly increased basal cardiomyocyte proliferation even in absence of BMP7, whereas SMAD5 knockdown trended towards a reduction in cardiomyocyte proliferation. Importantly, SMAD5 knockdown abolished cardiomyocyte proliferation induced by BMP7 exposure, whereas SMAD1 silencing did not affect BMP7 mitogenic activity (**Fig. 6c**). Finally, SMAD9 knockdown did not affect the level of basal or BMP7-induced neonatal cardiomyocyte proliferation (**Fig. 6c**). We further confirmed the activation of SMAD5 in enriched cardiomyocyte cultures, as evidenced by the increase of its phosphorylation levels upon BMP7 stimulation (**Fig. 6d; Supplementary Fig. 10d**). Interestingly, our data showed higher nuclear localization of phosphorylated SMAD1/5/9 *in vivo* in cardiomyocytes localized in the infarct border zone of BMP7-treated mice compared to control mice (**Fig. 6e**).

These data suggest that BMP7 triggers neonatal cardiomyocyte proliferation by activating SMAD signalling *in vitro* as well as *in vivo* in adult cardiomyocytes following myocardial infarction. SMAD5 specifically transduces the mitogenic signal of BMP7 in cardiomyocytes, while SMAD1 acts as an endogenous mitogenic suppressor. Additionally, SMAD9 does not appear to play a significant role in mediating BMP7-induced cardiomyocyte proliferation, likely due to its very low expression levels in cardiac muscle cells (**Supplementary Fig. 10e**).

### Non-canonical ERK and AKT pathways mediate BMP7 mitogenic signal transduction

BMP-activated receptors have been shown to activate several non-canonical pathways, including those involving extracellular signal-regulated kinases (ERKs) and protein kinase B (AKT) (Zhang, 2017). To investigate the potential role of non-canonical cascades downstream to BMP7 signalling, we tested the activation of the active phospho-isoforms of ERK and AKT. We observed a transient activation of ERK and AKT in cardiac cell cultures after BMP7 stimulation (**Fig. 7a**). By removing stromal cells by immunomagnetic separation, we confirmed that BMP7 treatment increases ERK activation in cardiomyocyte-enriched cultures (**Supplementary Fig. 11**). Surprisingly, we could not observe an increase of AKT activation in cardiomyocyte-enriched cultures upon BMP7 treatment, likely because the absence of stromal cells leads to overactivation of AKT at basal levels (**Supplementary Fig. 11**), a phenomenon that deserves further investigations. Importantly, our *in vivo* analyses showed increased nuclear immunoreactivity for ERK and AKT in cardiomyocytes within the border zone of BMP7-treated compared to control post-infarcted mice (**Fig. 7b-c**).

**Figure 7.**
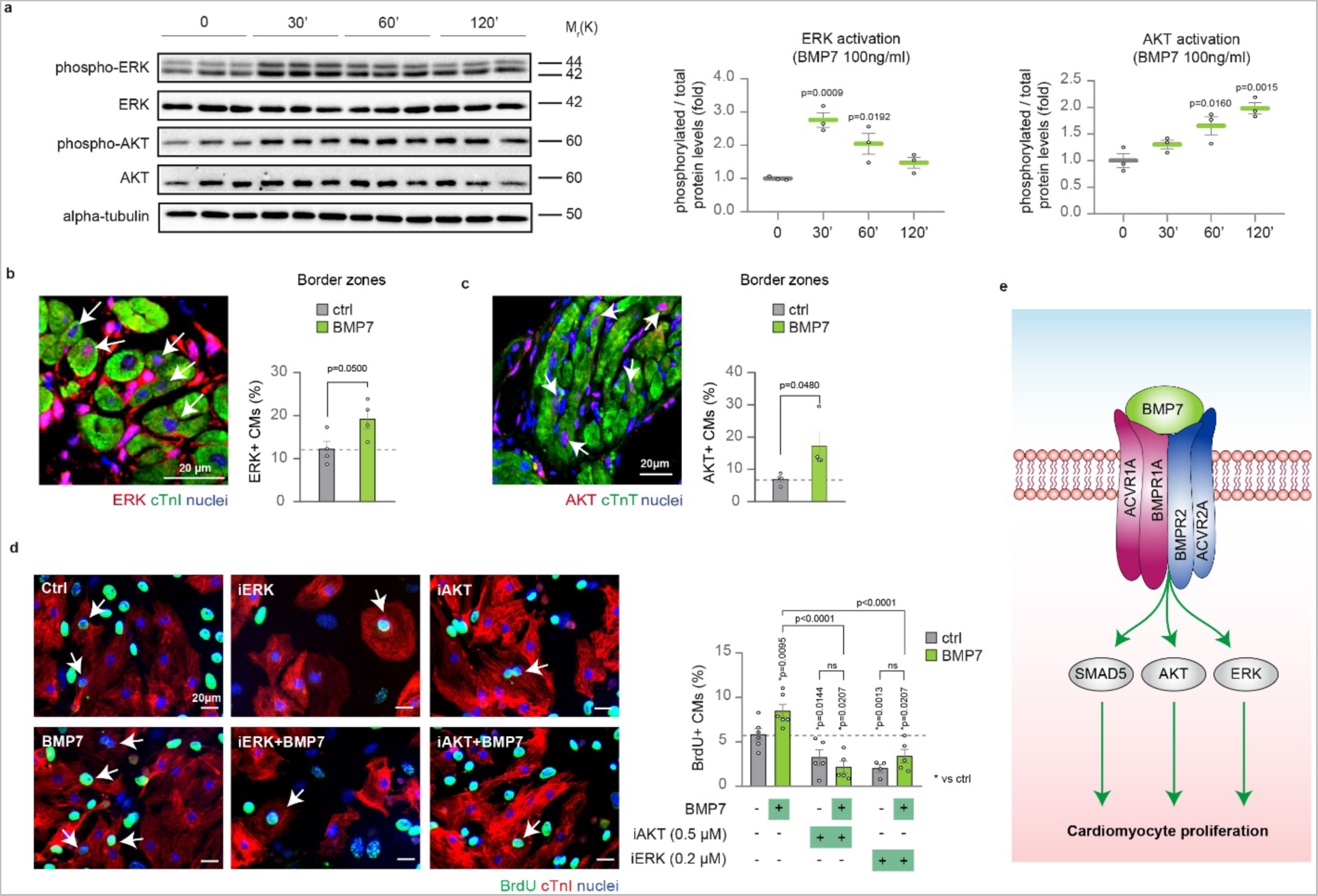
ERK and AKT non-canonical pathway take part in BMP7 mitogenic signal transduction. **(a)** Western Blot analysis of protein levels of phospho-ERK, ERK, phospho-AKT, AKT and alpha-tubulin in neonatal (postnatal day 1, P1) cardiac cultures following stimulation with 100 ng/ml BMP7 for 30, 60 and 120 minutes (n = 3 replicates per condition). Alpha-tubulin protein levels are provided as second loading control; (**b-c**) Immunofluorescence analysis for (**b**) ERK and cardiac Troponin I (cTnI) or (**c**) AKT and cardiac Troponin T (cTnT) in the border zones of heart sections 14 days post myocardial infarction, following daily injection of BMP7 or water as control (n = 8 mice; a total of 9318 and 4652 cardiomyocytes have been analysed in b and c, respectively). Cardiomyocytes with nuclear ERK or AKT immunoreactivity were counted manually within border zones in tissue sections; every dot represents a different heart (biological replicate), which in turn has been calculated by the analysis of 1-2 sections. Representative pictures are provided; arrows point at ERK-positive cardiomyocytes in panel **b** and at AKT-positive cardiomyocytes in panel **c**; scale bars 20 µm; (**d**) Quantification of BrdU incorporation in neonatal (postnatal day 1, P1) cardiomyocytes upon treatment with inhibitors of AKT (Akt inhibitor VIII 0.5 µM) or ERK (PD0325901 0.2 µM), with or without administration of BMP7 at 10 ng/mL for 48 hours (n = 8204 cardiomyocytes pooled from the analysis of 31 samples); cardiomyocytes were identified by cardiac Troponin I (cTnI) immunostaining. Representative pictures are provided; arrows point at proliferating cardiomyocytes; scale bars 20 µm; (**e**) BMP7 signal transduction model triggering cardiomyocyte proliferation, showing the involvement of canonical SMAD5 and non-canonical ERK and AKT. The values in **a**-**d** are presented as mean (error bars show s.e.m.), statistical significance was determined using one way ANOVA followed by Sidak’s test in **a, d** (comparison between pairs of treatments), and using two- sided Student’s t-test in **b** and **c**.

By using selective ERK and AKT inhibitors, we evaluated whether these pathways play a role in BMP7-induced cardiomyocyte proliferation. Inhibition of AKT and ERK reduced basal cardiomyocyte proliferation (**Fig. 7d**), which is consistent with previous observations (D’Uva et al., 2015a). Importantly, BMP7 was unable to enhance cardiomyocyte proliferation when activation of ERK or AKT was blocked (**Fig. 7d**). Overall, these data suggest that the mitogenic activity of BMP7 in cardiomyocytes is mediated both by the canonical SMAD5 and the non-canonical ERK and AKT pathways (**Fig. 7e**).

## Discussion

Several mitogens have been shown to promote the proliferation of cardiomyocytes during prenatal development and researchers have explored the administration of these factors or activation of downstream mediators as a strategy to restore the mitogenic potential of these cells, which is almost completely lost during the early postnatal life in mammals (reviewed in (Bongiovanni et al., 2021; Cahill et al., 2017; Galdos et al., 2017; Hashimoto et al., 2018; Heallen et al., 2019; Sadek and Olson, 2020; Tzahor and Poss, 2017; Uygur and Lee, 2016)). In this study, we hypothesised that specific growth factors play a crucial role in supporting cardiomyocyte proliferation until the early neonatal period. Subsequently, the declining expression of these factors leads to the exit of cardiomyocytes from the cell cycle, thereby reducing the cardiac regenerative ability.

To identify potential regenerative growth factors, we selectively considered those with diminishing expression levels during early postnatal mouse cardiac development, coinciding with the exit of cardiomyocytes from the cell cycle. Our data show that 18 of 23 identified growth factors effectively promote the proliferation of neonatal cardiomyocytes. This finding strongly supports our hypothesis. Notably, some of these growth factors, such as NRG1b (Bersell et al., 2009; D’Uva et al., 2015a; Polizzotti et al., 2015), IL6 (Tang et al., 2018), IGF2 (Shen et al., 2020; Zacchigna et al., 2018b) and sRANKL (Zacchigna et al., 2018b) and IL1b (Palmer et al., 1995), have been previously reported to possess pro-regenerative properties.

Our study also led to the identification of novel growth factors exerting a pro-proliferative effect on cardiac muscle cells, including CXCL17, BMP7, GDF5 (BMP14), LGALS7 (GAL7), CCL3, IL1RA, IL17b, CXCL14, IL10, IGFBP, LGALS3 (GAL3), IL17f and GDNF. Although NPPB, CTGF, IL23a, FGF23 and ARTN did not significantly promote cardiomyocyte proliferation in our study, it is possible that higher dosages, longer treatments, or combined administrations with these factors may still have a proliferative effect. Moreover, our analyses of cell cycle progression revealed that among the analysed factors, BMP7, a member of the bone morphogenetic protein family, exhibited the most potent induction of cardiomyocyte cell division. Notably, knockdown of BMP7 resulted in reduced proliferation of neonatal cardiomyocytes, indicating that endogenous production of BMP7 sustains cardiomyocyte proliferation at the neonatal stage. Our data also suggest that at the neonatal stage cardiomyocytes express *Bmp7* more abundantly than other cardiac cell populations. Thus, BMP7 appears to be an autocrine proliferative factor for cardiomyocytes, whose expression levels dramatically decline during the early postnatal life in mammals.

BMP7 has been shown to play an anti-fibrotic and anti-inflammatory role in various tissues, including the heart (Cao et al., 2022; D’Uva and Tzahor, 2015; Elmadbouh and Singla, 2021; Jin et al., 2018; Merino et al., 2016; Salido-Medina et al., 2022; Tan et al., 2019; Tate et al., 2021; Zeisberg et al., 2007) [reviewed in (Aluganti Narasimhulu and Singla, 2020; Weiskirchen et al., 2009)]. Administration of BMP7 has shown significant reduction in scar size and improvement in cardiac function after myocardial infarction in rat models (Jin et al., 2018). The beneficial effects of BMP7 treatment in reducing cardiac fibrosis have also been observed in models of pressure overload (Merino et al., 2016; Zeisberg et al., 2007), diabetic cardiomyopathy (Elmadbouh and Singla, 2021; Tate et al., 2021) and other cardiac genetic pathologies (Tan et al., 2019) and heart failure models (Cao et al., 2022). Dampening of TGF-β pro-fibrotic signalling has been identified as the mechanism by which BMP7 counteracts myocardial fibrosis (Chen et al., 2016). Our results suggest that BMP7 exerts a direct pro-regenerative effect on cardiomyocytes in addition to its anti-inflammatory and anti-fibrotic activities, thus making it a potential therapeutic strategy for triggering cardiac regeneration.

Interestingly, BMP7 did not show significant changes in expression levels after myocardial infarction in neonatal or adult mice. In line with the rationale of our initial screening, these findings suggest that the proliferative and regenerative ability of the neonatal mammalian heart depends at least in part on factors driving the physiological prenatal proliferation of cardiomyocytes, which are highly expressed at neonatal stage and do not require further upregulation in response to cardiac injury. Interestingly, adult zebrafish has the ability to re-activate some of these factors, such as bmp7a (Wu et al., 2016), while adult mice do not (as observed in this study). These observations may partially explain why fish regenerate even in adulthood while mice cannot. Nevertheless, screening of growth factors upregulated in regenerating hearts could serve as an alternative strategy for the identification of additional cardiac regenerative factors.

The BMP7 gene is highly conserved among vertebrate species. The importance of BMP signalling in cardiac regeneration was previously documented in the zebrafish model. Activation of the BMP/Smad pathway occurs after cardiac injury and BMP inhibition prevents zebrafish cardiomyocyte regeneration (Wu et al., 2016). Our study identifies *bmp7a* as an endogenous factor that supports cardiomyocyte proliferation during zebrafish cardiac regeneration and demonstrates that increasing *bmp7* expression is sufficient to enhance cardiomyocyte cell cycling. Further research is required to determine the role of other endogenous BMP ligands in the spontaneous regeneration of cardiomyocytes following cardiac injury in the zebrafish model. Notably, *bmp2b* has been shown to be upregulated after cardiac injury in the zebrafish model and its overexpression has been demonstrated to enhance zebrafish cardiomyocyte proliferation (Wu et al., 2016). Interestingly, in the mouse model, we observed a trend towards a reduction of *Bmp2* during the early postnatal period (see **Fig. 1a**). BMP2 administration has been shown to induce neonatal cardiomyocyte proliferation *in vitro* and to reduce cardiomyocyte apoptosis after myocardial infarction in adult mice (Chakraborty et al., 2013; Ebelt et al., 2013; Izumi et al., 2001). However, its potential to trigger mammalian cardiac regeneration remains unexplored.

Another member of the BMP family, BMP14 (GDF5), also exhibited a decline in expression during the early postnatal period in one of the two datasets analysed (see **Fig. 1a**) and its administration promoted cardiomyocyte cell cycle activity (see **Fig. 2**, **3a-b**). Increased expression of BMP14 after myocardial infarction has been reported to stimulate cardiomyocyte survival in the mouse model (Zaidi et al., 2010). Therefore, future studies may investigate BMP14 administration as cardiac regenerative strategy. Our study also elucidated the molecular mechanism underlying the mitogenic action of BMP7, identifying the receptors and downstream canonical and non-canonical signalling pathways. We found that two type I receptors (BMPR1A-ACVR1) and two type II receptors (BMPR2-ACVR2A) are crucial for mediating the mitogenic effect of BMP7 on cardiomyocytes, suggesting that a tetramer transduces BMP7 mitogenic signalling. Additionally, our study revealed that the activation of the canonical SMAD pathway plays a pivotal role in BMP7-induced cardiomyocyte proliferation. Interestingly, we observed that the BMP7 mitogenic effect on neonatal cardiomyocytes is specifically mediated by the canonical SMAD5 signalling, whereas SMAD1 appears to exert a paradoxical opposite role. BMPs have also been reported to activate multiple non-canonical, SMAD-independent signalling cascades, including those involving extracellular signal-regulated kinases (ERK), and the protein kinase B (AKT) (Zhang, 2017). Our study demonstrated that the BMP7 proliferative effect on cardiomyocytes is elicited through the activation of ERK and AKT. While ERK appears to be directly activated in cardiomyocytes following BMP7 treatment, AKT activation seems indirect and dependent on the presence of stromal cells. Further investigation is needed to understand the molecular mechanism underlying the dependence of AKT activation on cardiac stromal cells.

Furthermore, our study revealed that BMP7 reduces the proliferation of cardiac stromal cells both *in vitro* and *in vivo* after myocardial infarction. This finding aligns with the previously reported anti-proliferative effect of BMP7 on lung fibroblasts (Sun et al., 2018). Given that fibroblasts proliferate and migrate into the spaces created by dying cardiomyocytes after myocardial infarction (Le Bras, 2018; Eschenhagen, 2018; Fu et al., 2018), our data suggest an additional mechanism for the previously reported anti-fibrotic effect exerted by BMP7 on the cardiac tissue after injury (Jin et al., 2018). However, further investigation is needed to understand how BMP7 may induce an opposite effect on the proliferative ability of different cardiac cell types.

In conclusion, our study supports the concept that a temporal decline in the abundance of specific growth factors contributes to the exit of cardiomyocytes from the cell cycle and the loss of cardiac regenerative ability in mammals. Among these factors, BMP7, which is predominantly produced by cardiomyocytes in an autocrine manner, appears to be a potent inducer of mammalian cardiomyocyte proliferation both *in vitro* and *in vivo*. The mitogenic and pro-regenerative activity of *bmp7* on cardiomyocytes was also observed in the zebrafish model, in which heart regeneration naturally occurs. Therefore, we propose that administering BMP7 may be a novel strategy to enhance the regeneration of endogenous cardiomyocytes.

## ACKNOWLEDGMENTS

The research leading to these results has received funding from the European Union - NextGenerationEU through the Italian Ministry of University and Research under PNRR - M4C2-I1.3 Project PE_00000019 "HEAL ITALIA" to Gabriele Matteo D’Uva CUP J33C22002920006. The views and opinions expressed are those of the authors only and do not necessarily reflect those of the European Union or the European Commission. Neither the European Union nor the European Commission can be held responsible for them. The work was also supported by European Union’s Horizon 2020 research and innovation programme under the ERA-NET on Cardiovascular Diseases (ERA-CVD) Co-fund action to Gabriele Matteo D’Uva and Eldad Tzahor (Grant Number: JCT2016-40-080) and by Fondazione Luisa Fanti Melloni to Gabriele Matteo D’Uva. The work reported in this publication was funded by the Italian Ministry of Health.

Eldad Tzahor was supported by the European Research Council (ERC AdG #788194), the European Union’s Horizon 2020 research and innovation programme (874764), and the Israel Science Foundation (ISF).

Gilbert Weidinger was supported by the Deutsche Forschungsgemeinschaft (DFG, German Research Foundation) Project-ID 251293561 – SFB 1149 (project C03), project ID 316249678 – SFB 1279 (project Z02), and project ID 450627322 – SFB 1506 (project C04).

Part of this work was carried out in ALEMBIC, an advanced microscopy laboratory established by IRCCS Ospedale San Raffaele and Università Vita-Salute San Raffaele, and in the core facility “confocal and multiphoton microscopy” of the Medical faculty of Ulm University. We thank Milena Pariali for technical assistance in sectioning of paraffin-embedded samples, and Desirè Zambroni for technical assistance in the *in vitro* immunofluorescence imaging.

## AUTHOR CONTRIBUTIONS

C.B. and G.D’U. designed the experiments. C.B. carried out most of the experiments and analyzed the data. H.B.L. performed myocardial infarction experiments on mouse models. D.P.P., S.R., M.B., generated zebrafish transgenic lines and performed cryoinjury experiments in zebrafish models. L.K. cloned zebrafish constructs and D.P.P and L.K. performed experiments on zebrafish embryos. I.D.B., S.D.P., F.S., C.M., S.B., F.P., and I.P. performed immunofluorescence, western blots and gene expression analysis. R.T. helped with immunofluorescence image acquisition and time-lapse imaging. D.R., M.M. helped with western blot analysis. E.T., G.W., S.H., and M.L. supervised the experiments done by their laboratory members, and G.D’U. supervised the entire project. C.B. and G.D’U. wrote the manuscript, with editing contributions from all authors.

## MATERIALS AND METHODS

### Animal studies

Experiments involving mouse models were approved by the Animal Care and Use Committee of the University of Bologna (Italy), Weizmann Institute of Science (Israel), Cogentech Mouse Genetics Facility (IFOM, Milan, Italy). Experiments involving zebrafish were approved by the state of Baden-Württemberg and animal care representatives of Ulm University.

### Mouse cardiomyocyte isolation and culture

Primary neonatal cardiomyocytes were extracted from the hearts of 0-day-old or 1-day-old (P0-P1) mice. Neonatal cardiac cells were isolated by enzymatic digestion with pancreatin (Sigma) and collagenase (Roche), as previously described (Pianca et al., 2019). Primary juvenile cardiomyocytes were extracted from hearts of 7-day-old (P7) mice, which were anterogradely perfused with digestion enzymes (collagenase (Roche), trypsin (Sigma), and protease (Sigma)) as previously described (Omatsu-Kanbe et al., 2018). Cells were then cultured in 0.1% gelatine-coated (Sigma) wells with DMEM/F12 (Aurogene) supplemented with 1% L-glutamine (Sigma), 1% sodium pyruvate (Life Technologies), 1% non-essential amino acids (Life Technologies), 1% penicillin and streptomycin (Euroclone), 5% horse serum (Invitrogen) and 10% FBS (Life Technologies) (hereafter referred to as ‘complete-medium’) at 37°C and 5% CO_2_. The cells were allowed to adhere for 48 hours in complete-medium (24 hours for knockdown analysis). Subsequently, the medium was replaced with an FBS-deprived complete-medium containing selected growth factors (ImmunoTools) at the concentration of 10 ng/ml (or at the indicated concentrations in the figure legends), dosages in accordance with the usual range for *in vitro* experiments (D’Uva et al., 2015b; Engel et al., 2005; Kubin et al., 2011; Palmer et al., 1995; Sun et al., 2014; Wodsedalek et al., 2019), for about 48 hours (for proliferation analyses) or 30, 60 and 120 minutes (for cell signalling analysis). For BrdU assays, BrdU (10 µM, B5002, Sigma) was introduced along with the treatments. For Western blot analysis, 24 hours of starvation in the serum-deprived medium were performed before cell treatment.

### Transient gene knockdown

To silence specific molecular targets, postnatal-day 1 (P1) cardiomyocytes were isolated through enzymatic digestion and cultured in a complete medium, as described above. SMARTpool siRNAs, targeting Bmpr1a (L- 040598-00-0005, Dharmacon), Acvr1 (L-042047-00-0005, Dharmacon), Acvr1b (L- 043507-00-0005, Dharmacon), Acvrl1 (L-043004-00-0005, Dharmacon), Bmpr2 (L- 040599-00-0005, Dharmacon), Acvr2a (L-040676-00-0005, Dharmacon), Smad1 (L- 055762-00-0005, Dharmacon), Smad5 (L-057015-01-0005, Dharmacon) and Smad9 (L-046344-01-0005, Dharmacon), were delivered to cardiomyocytes, 24 hours post seeding, through Lipofectamine™ 3000 Transfection Reagent (L3000008, Thermofisher), following the manufacturer’s protocol. Lipofectamine™ 3000 Reagent was used at the lowest recommended concentration. Gene knockdown was obtained following 48 hours of transfection at 37°C and 5% CO_2._ Subsequently, cells were lysed for RNA extraction, as described below, or the medium was replaced with an FBS-deprived complete-medium containing the selected growth factors (ImmunoTools) for 48 hours (for proliferation assays).

### Mouse cardiomyocyte and stromal cell separation

To separate cardiomyocytes and stromal cells in P1 and P7 hearts, we proceeded as follows. P1 hearts were subjected to enzymatic digestion, whereas P7 hearts were harvested and anterogradely perfused as described above. Cardiomyocytes were separated from stromal cells using the immuno- magnetic cell sorting MACS Neonatal Isolation System (130-100-825, Miltenyi biotech), following the manufacturer’s protocol. Enriched cardiomyocytes were then cultured in a complete medium. The validation of the enrichment procedure was previously demonstrated by the analysis of gene expression levels of markers specific for cardiomyocytes and stromal cells (fibroblasts and endothelial cells) (Pianca et al., 2022). To assess changes in gene expression levels cardiomyocytes were centrifuged, and the total RNA was extracted from the pellet as described below.

### Immunofluorescence on mouse cells and tissue sections

Cultured cells were fixed with 4% paraformaldehyde (PFA) solution (Sigma, diluted in PBS) for 20 minutes at 4°C or room temperature and then washed thrice in PBS. Heart sections (4 µm) underwent deparaffinization (by immersion in Toluene and rehydration by immersion in solutions with a decreasing concentration of Ethanol), and heat-induced antigen retrieval in EDTA buffer (Sigma) at pH 8.5 (for Ki67, phospho-SMAD1/5/9 and ERK2 immunostaining) or Sodium Citrate buffer (10mM Sodium Citrate, 0.05% Tween 20) at pH 6 (for Aurora B kinase and BrdU immunostaining), followed by gradual chilling. Then heart sections and cultured cells were processed in the following manner. Samples were permeabilized with 0.5% Triton-X100 (Sigma) in PBS for 5 minutes at room temperature and the non-specific binding of the antibodies was prevented by applying a blocking solution (PBS supplemented with 5% BSA (Sigma) or 5% goat serum (for Aurora B kinase immunostaining in heart sections) and 0.1% Triton-X100) for 1 hour at room temperature. For the BrdU staining protocol on cultured cells, a DNA hydrolysis step between the permeabilization and the blocking step was performed by the addition of 2M HCL (Sigma) for 30 minutes at 37°C, followed by 3 washes in PBS. For the BrdU staining protocol on tissue section a DNA hydrolysis step directly after antigen retrieval was performed by the addition of 2M HCL (Sigma) in PBS for 30 minutes at 37°C, followed by a step of HCL neutralization with 0.1 M Borate Buffer pH 8.5 for 10 minutes at room temperature, 3 washes in PBS and incubation with blocking solution (PBS supplemented with 20% horse serum and 0.5% Triton-X100). Then samples were incubated overnight at 4°C with primary antibodies diluted in PBS, supplemented with 3% BSA or 3% goat serum (for Aurora B kinase, phospho- SMAD1/5/9 and ERK2 analysis in heart sections) and 0.1% Triton-X100. For the BrdU staining protocol primary antibodies diluted in PBS, supplemented with 2% horse serum and 0.5% Triton-X100 were used. Anti-Troponin T (cTnT) (1:400, ab33589, Abcam) and anti-Troponin I (cTnI) (1:400, ab47003, Abcam) antibodies were used to identify cardiomyocytes. Anti-KI67 (1:50, ab16667, Abcam), anti-BrdU (1:40, G3G4, DSHB), and anti-Aurora B kinase (1:50, 611082, BD Transduction Laboratories) antibodies were used to analyse cell-cycle activity, DNA synthesis and cytokinesis, respectively. Anti-phospho SMAD 1/5/9 (1:500, 13820, Cell Signaling) and anti-ERK2 (1:50, sc-1647, Santa Cruz) were used to analyse BMP7 downstream signalling after myocardial infarction.

After primary antibody incubation, 3 washes in PBS were performed and the samples were incubated for 1 hour at room temperature with fluorescent secondary antibodies, diluted (1:200) in PBS supplemented with 1% BSA, 1% goat serum (for Aurora B kinase, phospho-SMAD1/5/9 and ERK2 analysis in heart sections) or 1% horse serum (for BrdU analysis in heart sections) and 0.1% Triton-X100. For the BrdU staining protocol tissue sections were incubated for 90 minutes at room temperature with fluorescent secondary antibodies, diluted (1:200) in PBS supplemented with 2% horse serum. The following secondary antibodies were used: anti-mouse AlexaFluor 488 (115-545-003, Jackson), anti-rabbit AlexaFluor 488 (111-545-003, Jackson), anti-mouse Cy3 (115-165-003, Jackson), anti-rabbit 594 (AlexaFluor 111-585-003, Jackson), anti-rabbit Cy3 (111-165-003, Jackson). After 3 washes in PBS, DAPI (4’,6-diamidino-2-phenylindole dihydrochloride, Sigma), diluted at 1 µg/ml in PBS, was applied for 10 minutes at room temperature for nuclei visualization. 15 minutes of incubation were applied for the BrdU staining protocol on cultured cells. Samples were then washed 2 more times in PBS. Cells in the culture plates were imaged with the following widefield microscopies: Zeiss (Axio Observer A1), ArrayScan XTI (ThermoFisher), and Eclipse Ti2 (Nikon) at x20 magnification. Slides were mounted with an antifade solution (Vectorlabs), covered with a coverslip, sealed with nail polish and imaged at an Olympus VS200 slide scanner at x20 magnification. Z-stack images were acquired at Eclipse Ti2 (Nikon) with CrestOptics DeepSIM at x60 magnification. Images were analysed with Photoshop or Image J software.

KI67, Aurora B kinase, BrdU, phosphor-SMAD1/5/9 and ERK2 positive cardiomyocytes in tissue sections were counted manually within infarcted plus border zones or remote zones. For each heart an average value was calculated from the analysis of 1 to 3 sections per heart and then inserted in a boxplot. KI67 positive stromal cells in tissue sections were counted manually within infarcted plus border zones from 12 randomly selected circular areas; for each heart an average value was calculated from the analysis of 2 to 3 sections per heart and then inserted in a boxplot.

### Protein extraction and Western Blotting

Western blotting was performed with the SDS–PAGE Electrophoresis System. Proteins were extracted from 400,000 to 500,000 cells with RIPA buffer with the addition of proteinase inhibitor (Sigma) and phosphatase inhibitors (Sigma). Then, 20-25 µg protein extracts were resolved by sodium dodecyl sulfate (SDS)-polyacrylamide gel electrophoresis and transferred to a nitrocellulose membrane (AmershamTM ProtranTM Premium 0.45 µm 300 mm x 4 m). The membrane was blocked for 1 hours using TBS-T (0.1% Tween-20) supplemented by 5% BSA (Sigma), and incubated overnight at 4°C with the following primary antibodies: anti-SMAD 1 (1:1000, 6944, Cell Signaling), anti-phospho SMAD 1/5/9 (1:1000, 13820, Cell Signaling), anti-ERK2 (1:1000, sc-1647, Santa Cruz), anti-phospho-ERK (1:2500, M8159, Sigma), anti-AKT (1:1000, 9272, Cell Signaling), anti-phospho-AKT (1:1000, 9271, Cell Signaling), anti-GAPDH (1:1000, 2118, Cell Signaling,) and anti-α-tubulin (1:4000, T5168, Sigma). For protein detection, the membrane was incubated with anti-rabbit or anti-mouse horseradish peroxidase-labelled secondary antibody (Dako EnVision+ System- HRP Labelled Polymer) followed by a chemiluminescent reaction (Clarity Western ECL Substrate, Bio-Rad).

Signals and images were acquired by ChemiDoc™ XRS 2015 (Bio-Rad Laboratories), and densitometric analysis were performed using Image Lab software (version 5.2.1; Bio-Rad Laboratories).

### Transcriptional analysis

Total RNA extraction was performed with the NucleoSpin RNA II kit (Macherey Nagel) according to the manufacturer’s protocol. RNA quantification and quality check were performed using a Nanodrop spectrophotometer (N1000, Thermo). RNA was reverse transcribed to double-stranded cDNA using SuperScript™ VILO™ cDNA Synthesis Kit (Invitrogen) according to the manufacturer’s protocol. Real-Time (rt)-PCR was performed using Fast SYBR Green PCR Master Mix (Applied Biosystems) on a QuantStudio 6 Flex instrument (Applied Biosystems) and a QuantStudio 5 Flex instrument (Applied Biosystems). Oligonucleotide sequences of genes analysed in this study namely *Bmp7*, *Bmpr1a*, *Bmpr2*, *Acvr2a* and *Acvr1*, are listed in **Supplementary Table 2**. Relative quantification was performed using the *Hprt1* gene as a loading control. DDCT was calculated and data of each gene were analysed using a 2^-DDCT^ method and reported as mean fold change.

### Time-lapse imaging

Following cell isolation, postnatal day 1 (P1) cardiomyocytes were seeded at 35,000 cells per well in a 96-well plate and left to adhere for 48 hours. Then, the complete medium was replaced with serum-free medium supplemented with 10 nM TMRE (tetramethylrhodamine ethyl ester, Sigma), a fluorescent mitochondrial dye used to label cardiomyocytes (Hattori et al., 2010). After 20 minutes of incubation at 37°C, the medium containing TMRE was removed, and cardiomyocytes were treated with and without BMP7 (ImmunoTools) at 10 ng/ml in FBS-deprived complete medium. Live cell imaging was performed using a widefield fluorescent microscope (Nikon Eclipse TI2). Time-lapse imaging was carried out for 16 hours and images were acquired at x40 magnification every 15 minutes.

### Myocardial infarction in the mouse model

Myocardial infarction in the adult stage was induced by ligation of the left anterior descending coronary artery as previously described (D’Uva et al., 2015). Female 3-months old C57BL/6JRccHsd mice were sedated with isoflurane (Abbott Laboratories) and artificially ventilated following tracheal intubation. Lateral thoracotomy at the fourth intercostal space was performed by blunt dissection of the intercostal muscles following skin incision. After the ligation of the left anterior descending coronary artery, thoracic wall incisions were sutured with 6.0 non-absorbable silk sutures, and the skin wound was closed using glue. Mice were then warmed for several minutes until recovery.

### *In vivo* drug delivery in the mouse model

Two days after myocardial infarction, Recombinant Human BMP7 (Peprotech) was delivered on a daily basis with a dose of 100 ug/kg/day dissolved in water for 12 days, through IV (intravenous) injection every 3 days and IP (intraperitoneal) injection for the rest of the time up to 14 days after myocardial infarction. The administration method and the selected dose of the growth factor are similar to previous reports (Bersell et al., 2009). Infarcted control mice were subjected to daily injections of water for 12 days. At the end of the treatment, mice were sacrificed, and hearts were collected for analysis of cardiac cell proliferation (KI67, Aurora B kinase and BrdU staining) or BMP7 downstream pathways (phospho- SMAD1/5/9, ERK2 and AKT staining). For the BrdU staining protocol 50 mg kg^−1^ BrdU were injected at day 3, 5 and 7 post-myocardial infarction.

### Rescue of zebrafish *bmp7a* (*snailhouse*) mutants

We previously reported that *bmp7* expression is induced after ventricular cryoinjury of adult zebrafish hearts (Wu et al., 2016). Please note that the transcript termed “*bmp7*” in our previous publication (Wu et al., 2016) has in the meantime been re-named *bmp7a* by the zebrafish nomenclature committee. We used the following previously identified *bmp7a* loss-of-function allele in the present study: *snh^ty68a^*(Schmid et al., 2000).

To test the efficacy of zebrafish *bmp7a* and *bmp7b* to rescue the early developmental defects of *bmp7a* homozygous mutants, the *bmp7a* and *bmp7b* coding sequences (ZDB-GENE-000208-25 and ZDB-GENE-060929-328 respectively) were cloned into the pCS2P+ vector. Capped sense RNA was synthesized from linearized plasmids using mMessage mMachine kit (Invitrogen) and RNA was purified using RNeasy Mini kit (Qiagen). 300 pg of *bmp7a* or *bmp7b* mRNA was injected into embryos derived from incrosses of *bmp7a*^ty68a^ heterozygous carriers at the 1 cell stage. Injected embryos were allowed to grow until 3 dpf (days post fertilization), at which stage all non-rescued homozygous mutant embryos would have died. Genomic DNA was isolated from single embryos using NaOH, diluted 1:6 in nuclease free H_2_O and 5 µl were used for Kompetitive allele specific PCR (KASP) genotyping (LGC Biosearch Technology) (**Supplementary Table 3**). The 2X KASP Master mix and KASP Assay primer mix were designed and provided by LGC Biosearch Technology (KASP by Design) using TACTCTTATGAACCCGCGTACACGACCCCGGGACCCCCGCTGGTGACCCAGCAGGACAGTCGCTTTCTCAGTGATGCCGACATGG**[T/G]**GATGAGCTTTGC GAATACAGGTGAGCGTCTTATGAAATTCACCGCATATCATAATTGTTGTTAGGATGAATCAACAGATTGTTTTTGCTCCATT for each allele in which the FAM fluorophore (T) corresponds to the wild type allele and the HEX fluorophore (G) corresponds to the mutated one. The rescue efficiency was calculated by dividing the % mutant genotype by the % expected mutant genotype after complete rescue.

To generate *bmp7a* homozygous mutants for adult experiments, embryos derived from incrosses of *snh^ty68a^* heterozygous carriers were injected with 300 pg of *bmp7a* mRNA, raised to adulthood and homozygous fish were identified by sequencing. To this end, a fragment of the *bmp7a* locus was amplified by PCR using the primers 5’ TTGTGTACTCTTATGAACCCGCGTA 3’ (sense) and 5’ GGAGCAAAAACAATCTGTTGATTCA 3’ (antisense) 3’ and PCR products were sequenced using the sense primer. The *ty68a* allele contains a single T-to-G point mutation: the wild-type sequence is GTG (coding for Valine), the mutant sequence is GGG (coding for Glycine). Adult tanks contained on average 12% homozygous mutant fish; thus we were able to rescue about half of the expected homozygous mutant embryos (25% of the total).

### Phenotypic comparison between *bmp7a* and *bmp7b* overexpression models

To compare the biological activity of zebrafish *bmp7a* and *bmp7b* when overexpressed, 300 pg of *bmp7a* or *bmp7b* mRNA was injected into zebrafish embryos derived from wild-type outcrosses at the 1 cell stage. Phenotype analysis was performed at 24 hpf (hours post fertilisation) and classified into the V1-V4 ventralisation classes scheme according to previously published data (Kishimoto et al., 1997).

### Generation of *hsp70l*:bmp7b, *myl7*:eGFP^af5Tg^ transgenic zebrafish line

The following transgenic line for heat-shock-inducible overexpression of zebrafish *bmp7* and concomitant labelling of cardiomyocytes by eGFP was created for this publication: *hsp70l*:bmp7b, *myl7*:eGFP^af5Tg^. To this end the coding sequence of *bmp7b* (ZFIN-ID: ZDB-GENE-060929-328) was amplified from cDNA and cloned into pENTR/D-TOPO (Thermo Fisher). The transgenesis construct was generated by Multiway Gateway (Thermo Fisher) cloning using pENTR-bmp7b, as well as p5E- *hsp70l*, p3E-pA, and pDestTol2CG2 from the Tol2 kit (Kwan et al., 2007). AB wild type embryos were injected with *hsp70l*:bmp7b, *myl7*:eGFP plasmid and Tol2 transposase mRNA, screened for *myl7*:GFP expression, and raised to adulthood. Mosaic F0 adults were crossed with AB wild type fish to obtain heterozygous tg(*hsp70l*:bmp7b, *myl7*:GFP) zebrafish.

### Zebrafish heart cryoinjuries, EdU injection and heat-shocks

Cryoinjuries were performed as previously described on zebrafish of 4-6 months of age (Wu et al., 2016), except that a liquid nitrogen-cooled copper filament of 0.3-mm diameter was used instead of dry ice. In experiments involving *bmp7a* loss-of-function, fish were injected intraperitoneally with 10 μl of 10 mM 5-ethynyl-2-deoxyuridine (EdU) in PBS once daily for 3 days, starting 4 days after cryoinjury, and cycling of cardiomyocytes (detected by anti-MHC immunofluorescence) was evaluated at 7 days post injury. For heat-shock-inducible overexpression of *bmp7b*, *hsp70l*:bmp7b, *myl7*:eGFP^af5Tg^ transgenic fish were heat-shocked once daily for 6 days, starting at 1 day post cryoinjury (dpi), and analysed 5 hours after the end of the last heat shock at 7 dpi. To apply heat-shocks, water containing fish was heated to 37°C for 1 hour, after which the water temperature was reduced back to 27°C within 15 min. Wild-type negative control fish were heat-shocked as well.

### Immunofluorescence on zebrafish tissue sections

Zebrafish hearts were fixed in 4% paraformaldehyde (PFA) (in phosphate buffer with 4% sucrose) at room temperature for 1 hour, washed three times for 10 minutes in 4% sucrose/phosphate buffer and equilibrated in 30% sucrose/phosphate buffer overnight at 4°C. Hearts were embedded and cryosectioned into 10-μm sections. Sections were equally distributed onto six serial slides so that each slide contained sections representing all areas of the ventricle. Immunostainings were performed as previously described (Schnabel et al., 2011). The following primary antibodies were used: anti-PCNA (1:1000, M0879, RRID:AB_2160651, Dako), anti-MF20 (which detects sarcomeric Myosin heavy chain – MHC; 1:50, MF20, RRID:AB_2147781, Developmental Studies Hybridoma Bank), anti-Myl7 (1:300, GTX128346, RRID:AB_2885759, Genetex). Secondary antibodies conjugated to Alexa 488, 555 or 633 (Invitrogen) were used at a dilution of 1:1000. Nuclei were shown by DAPI (40,60- diamidino-2-phenylindole) staining.

For EdU detection, cryosections were prepared the same way as for immunofluorescence staining. Slides were washed twice in 3% BSA/PBS for 5 minutes, once in 0.5% Triton X-100/PBS for 20 minutes, and twice in 3% BSA/PBS for 3 minutes each. Slides were incubated in the dark with the reaction cocktail (EdU- Imaging kit, baseclick GmbH) as per manufacturer’s protocol for 30 minutes. Slides were washed for 3 minutes in 3% BSA/PBS and subjected to the immunofluorescence protocol to detect MHC (MF20) as described above.

All images of immunofluorescence staining are single optical planes acquired using 20x magnification with a Zeiss AxioObserver 7 equipped with an Apotome or with Leica Sp5 or Sp8 confocal microscopes. Quantifications of the fraction of EdU or PCNA positive cardiomyocytes were performed on 2 to 3 sections with the biggest wounds per heart and in cardiomyocytes located within 150 µm from the wound border. EdU+ or PCNA+ cardiomyocytes were counted manually within this border zone area. The number of cardiomyocytes in the wound border zone was estimated by multiplying the wound border area size (measured using ImageJ) with the average density of cardiomyocytes in three separate regions of interest (size: 100 µm^2^) within the border zone area, which was determined by counting cardiomyocyte nuclei within the regions of interest. For each heart the average value was calculated from the analysis of 2-3 sections per heart.

#### Statistical analysis

Statistical analyses were performed using GraphPad Prism 8 software. Whenever normality could be assumed, the two-sided Student’s t-test or analysis of variance (ANOVA) followed by Sidak’s, Tukey’s and Dunnet’s test were used to compare group means, as specified in the figure legends. P value<0.05 was considered to represent a statistically significant difference. In all panels, numerical data are expressed as mean+standard error of the mean (s.e.m.). For zebrafish phenotypic analysis Chi-squared test has been used.

## Data availability

RNA-sequencing data analysed in this study are available as Supplementary information within the original articles (Haubner et al., 2012; Talman et al., 2018).

## References

Aluganti Narasimhulu, C., and Singla, D.K. (2020). The Role of Bone Morphogenetic Protein 7 (BMP-7) in Inflammation in Heart Diseases. Cells 9, 1–30.

Benjamin, E.J., Muntner, P., Alonso, A., Bittencourt, M.S., Callaway, C.W., Carson, A.P., Chamberlain, A.M., Chang, A.R., Cheng, S., Das, S.R., et al. (2019). Heart Disease and Stroke Statistics-2019 Update: A Report From the American Heart Association.

Bergmann, O., Bhardwaj, R.D., Bernard, S., Zdunek, S., Barnabé-Heider, F., Walsh, S., Zupicich, J., Alkass, K., Buchholz, B.A., Druid, H., et al. (2009). Evidence for cardiomyocyte renewal in humans. Science 324, 98–102.

van Berlo, J.H., and Molkentin, J.D. (2014). An emerging consensus on cardiac regeneration. Nat. Med. 20, 1386–1393.

Bersell, K., Arab, S., Haring, B., and Kuhn, B. (2009). Neuregulin1/ErbB4 signaling induces cardiomyocyte proliferation and repair of heart injury. Cell 138, 257–270.

Bongiovanni, C., Sacchi, F., Da Pra, S., Pantano, E., Miano, C., Morelli, M.B., and D’Uva, G. (2021). Reawakening the Intrinsic Cardiac Regenerative Potential: Molecular Strategies to Boost Dedifferentiation and Proliferation of Endogenous Cardiomyocytes. Front. Cardiovasc. Med. 8, 750604.

Le Bras, A. (2018). Dynamics of fibroblast activation in the infarcted heart. Nat. Rev. Cardiol. 15, 379.

Bryant, D.M., O’Meara, C.C., Ho, N.N., Gannon, J., Cai, L., and Lee, R.T. (2015). A systematic analysis of neonatal mouse heart regeneration after apical resection. J. Mol. Cell. Cardiol. 79, 315–318.

Cahill, T.J., Choudhury, R.P., and Riley, P.R. (2017). Heart regeneration and repair after myocardial infarction: translational opportunities for novel therapeutics. Nat. Rev. Drug Discov. 16, 699–717.

Cao, Y., Wang, Y., Zhou, Z., Pan, C., Jiang, L., Zhou, Z., Meng, Y., Charugundla, S., Li, T., Allayee, H., et al. (2022). Liver-heart cross-talk mediated by coagulation factor XI protects against heart failure. Science 377, 1399–1406.

Chakraborty, S., Sengupta, A., and Yutzey, K.E. (2013). Tbx20 promotes cardiomyocyte proliferation and persistence of fetal characteristics in adult mouse hearts. J. Mol. Cell. Cardiol. 62, 203–213.

Chen, X., Xu, J., Jiang, B., and Liu, D. (2016). Bone Morphogenetic Protein-7 Antagonizes Myocardial Fibrosis Induced by Atrial Fibrillation by Restraining Transforming Growth Factor-β (TGF-β)/Smads Signaling. Med. Sci. Monit. 22, 3457–3468.

D’Uva, G., and Tzahor, E. (2015). The key roles of ERBB2 in cardiac regeneration. Cell Cycle 14, 2383–2384.

D’Uva, G., Aharonov, A., Lauriola, M., Kain, D., Yahalom-Ronen, Y., Carvalho, S., Weisinger, K., Bassat, E., Rajchman, D., Yifa, O., et al. (2015a). ERBB2 triggers mammalian heart regeneration by promoting cardiomyocyte dedifferentiation and proliferation. Nat. Cell Biol. 17, 627–638.

D’Uva, G., Aharonov, A., Lauriola, M., Kain, D., Yahalom-Ronen, Y., Carvalho, S., Weisinger, K., Bassat, E., Rajchman, D., Yifa, O., et al. (2015b). ERBB2 triggers mammalian heart regeneration by promoting cardiomyocyte dedifferentiation and proliferation. Nat. Cell Biol. 17, 627–638.

Derks, W., and Bergmann, O. (2020). Polyploidy in Cardiomyocytes: Roadblock to Heart Regeneration? Circ. Res. 126, 552–565.

Dong, X.-R., Wan, S.-M., Zhou, J.-J., Nie, C.-H., Chen, Y.-L., Diao, J.-H., and Gao, Z.-X. (2022). Functional Differentiation of BMP7 Genes in Zebrafish: bmp7a for Dorsal-Ventral Pattern and bmp7b for Melanin Synthesis and Eye Development. Front. Cell Dev. Biol. 10, 838721.

Drenckhahn, J.-D., Schwarz, Q.P., Gray, S., Laskowski, A., Kiriazis, H., Ming, Z., Harvey, R.P., Du, X.-J., Thorburn, D.R., and Cox, T.C. (2008). Compensatory growth of healthy cardiac cells in the presence of diseased cells restores tissue homeostasis during heart development. Dev. Cell 15, 521–533.

Ebelt, H., Hillebrand, I., Arlt, S., Zhang, Y., Kostin, S., Neuhaus, H., Müller-Werdan, U., Schwarz, E., Werdan, K., and Braun, T. (2013). Treatment with bone morphogenetic protein 2 limits infarct size after myocardial infarction in mice. Shock 39, 353–360.

Elmadbouh, I., and Singla, D.K. (2021). BMP-7 Attenuates Inflammation-Induced Pyroptosis and Improves Cardiac Repair in Diabetic Cardiomyopathy. Cells 10.

Engel, F.B., Schebesta, M., Duong, M.T., Lu, G., Ren, S., Madwed, J.B., Jiang, H., Wang, Y., and Keating, M.T. (2005). p38 MAP kinase inhibition enables proliferation of adult mammalian cardiomyocytes. Genes Dev. 19, 1175–1187.

Engel, F.B., Hsieh, P.C.H., Lee, R.T., and Keating, M.T. (2006). FGF1/p38 MAP kinase inhibitor therapy induces cardiomyocyte mitosis, reduces scarring, and rescues function after myocardial infarction. Proc. Natl. Acad. Sci. U. S. A. 103, 15546– 15551.

Eschenhagen, T. (2018). A new concept of fibroblast dynamics in post-myocardial infarction remodeling. J. Clin. Invest. 128, 1731–1733.

Eschenhagen, T., Bolli, R., Braun, T., Field, L.J., Fleischmann, B.K., Frisén, J., Giacca, M., Hare, J.M., Houser, S., Lee, R.T., et al. (2017). Cardiomyocyte Regeneration: A Consensus Statement. Circulation 136, 680–686.

Fu, X., Khalil, H., Kanisicak, O., Boyer, J.G., Vagnozzi, R.J., Maliken, B.D., Sargent, M.A., Prasad, V., Valiente-Alandi, I., Blaxall, B.C., et al. (2018). Specialized fibroblast differentiated states underlie scar formation in the infarcted mouse heart. J. Clin. Invest. 128, 2127–2143.

Galdos, F.X., Guo, Y., Paige, S.L., VanDusen, N.J., Wu, S.M., and Pu, W.T. (2017). Cardiac Regeneration: Lessons From Development. Circ. Res. 120, 941–959.

González-Gómez, P., Crecente-Campo, J., Zahonero, C., de la Fuente, M., Hernández-Laín, A., Mira, H., Sánchez-Gómez, P., and Garcia-Fuentes, M. (2015). Controlled release microspheres loaded with BMP7 suppress primary tumors from human glioblastoma. Oncotarget 6, 10950–10963.

Hashimoto, H., Yuasa, S., Tabata, H., Tohyama, S., Hayashiji, N., Hattori, F., Muraoka, N., Egashira, T., Okata, S., Yae, K., et al. (2014). Time-lapse imaging of cell cycle dynamics during development in living cardiomyocyte. J. Mol. Cell. Cardiol. 72, 241–249.

Hashimoto, H., Olson, E.N., and Bassel-Duby, R. (2018). Therapeutic approaches for cardiac regeneration and repair. Nat. Rev. Cardiol. 15, 585–600.

Hattori, F., Chen, H., Yamashita, H., Tohyama, S., Satoh, Y.-S., Yuasa, S., Li, W., Yamakawa, H., Tanaka, T., Onitsuka, T., et al. (2010). Nongenetic method for purifying stem cell-derived cardiomyocytes. Nat. Methods 7, 61–66.

Haubner, B.J., Adamowicz-Brice, M., Khadayate, S., Tiefenthaler, V., Metzler, B., Aitman, T., and Penninger, J.M. (2012). Complete cardiac regeneration in a mouse model of myocardial infarction. Aging (Albany. NY). 4, 966–977.

Heallen, T.R., Kadow, Z.A., Kim, J.H., Wang, J., and Martin, J.F. (2019). Stimulating Cardiogenesis as a Treatment for Heart Failure. Circ. Res. 124, 1647–1657.

Heldin, C.-H., and Moustakas, A. (2016). Signaling Receptors for TGF-β Family Members. Cold Spring Harb. Perspect. Biol. 8.

Hirose, K., Payumo, A.Y., Cutie, S., Hoang, A., Zhang, H., Guyot, R., Lunn, D., Bigley, R.B., Yu, H., Wang, J., et al. (2019). Evidence for hormonal control of heart regenerative capacity during endothermy acquisition. Science (80-.). 364, 184–188.

Izumi, M., Fujio, Y., Kunisada, K., Negoro, S., Tone, E., Funamoto, M., Osugi, T., Oshima, Y., Nakaoka, Y., Kishimoto, T., et al. (2001). Bone morphogenetic protein-2 inhibits serum deprivation-induced apoptosis of neonatal cardiac myocytes through activation of the Smad1 pathway. J. Biol. Chem. 276, 31133–31141.

Jin, Y., Cheng, X., Lu, J., and Li, X. (2018). Exogenous BMP-7 Facilitates the Recovery of Cardiac Function after Acute Myocardial Infarction through Counteracting TGF-β1 Signaling Pathway. Tohoku J. Exp. Med. 244, 1–6.

Jopling, C., Sleep, E., Raya, M., Martí, M., Raya, A., Belmonte, J.C.I., and Izpisúa Belmonte, J.C. (2010). Zebrafish heart regeneration occurs by cardiomyocyte dedifferentiation and proliferation. Nature 464, 606–609.

Kikuchi, K., Holdway, J.E., Werdich, A.A., Anderson, R.M., Fang, Y., Egnaczyk, G.F., Evans, T., Macrae, C. a, Stainier, D.Y.R., and Poss, K.D. (2010). Primary contribution to zebrafish heart regeneration by gata4(+) cardiomyocytes. Nature 464, 601–605.

Kishimoto, Y., Lee, K.H., Zon, L., Hammerschmidt, M., and Schulte-Merker, S. (1997). The molecular nature of zebrafish swirl: BMP2 function is essential during early dorsoventral patterning. Development 124, 4457–4466.

Kubin, T., Pöling, J., Kostin, S., Gajawada, P., Hein, S., Rees, W., Wietelmann, A., Tanaka, M., Lörchner, H., Schimanski, S., et al. (2011). Oncostatin M is a major mediator of cardiomyocyte dedifferentiation and remodeling. Cell Stem Cell 9, 420– 432.

Kwan, K.M., Fujimoto, E., Grabher, C., Mangum, B.D., Hardy, M.E., Campbell, D.S., Parant, J.M., Yost, H.J., Kanki, J.P., and Chien, C.-B. (2007). The Tol2kit: a multisite gateway-based construction kit for Tol2 transposon transgenesis constructs. Dev. Dyn. 236, 3088–3099.

Lavery, K., Swain, P., Falb, D., and Alaoui-Ismaili, M.H. (2008). BMP-2/4 and BMP- 6/7 differentially utilize cell surface receptors to induce osteoblastic differentiation of human bone marrow-derived mesenchymal stem cells. J. Biol. Chem. 283, 20948–20958.

Li, F., Wang, X., Capasso, J.M., and Gerdes, A.M. (1996). Rapid transition of cardiac myocytes from hyperplasia to hypertrophy during postnatal development. J. Mol. Cell. Cardiol. 28, 1737–1746.

Li, Y., Feng, J., Song, S., Li, H., Yang, H., Zhou, B., Li, Y., Yue, Z., Lian, H., Liu, L., et al. (2020a). gp130 Controls Cardiomyocyte Proliferation and Heart Regeneration. Circulation 142, 967–982.

Li, Z., Hu, S., Huang, K., Su, T., Cores, J., and Cheng, K. (2020b). Targeted anti-IL- 1β platelet microparticles for cardiac detoxing and repair. Sci. Adv. 6, eaay0589.

Loomans, H.A., and Andl, C.D. (2016). Activin receptor-like kinases: A diverse family playing an important role in cancer. Am. J. Cancer Res. 6, 2431–2447.

Massagué, J., Seoane, J., and Wotton, D. (2005). Smad transcription factors. Genes Dev. 19, 2783–2810.

Merino, D., Villar, A. V, García, R., Tramullas, M., Ruiz, L., Ribas, C., Cabezudo, S., Nistal, J.F., and Hurlé, M.A. (2016). BMP-7 attenuates left ventricular remodelling under pressure overload and facilitates reverse remodelling and functional recovery. Cardiovasc. Res. 110, 331–345.

Miyazono, K., Maeda, S., and Imamura, T. (2005). BMP receptor signaling: Transcriptional targets, regulation of signals, and signaling cross-talk. Cytokine Growth Factor Rev.

Novoyatleva, T., Diehl, F., van Amerongen, M.J., Patra, C., Ferrazzi, F., Bellazzi, R., and Engel, F.B. (2010). TWEAK is a positive regulator of cardiomyocyte proliferation. Cardiovasc. Res. 85, 681–690.

Omatsu-Kanbe, M., Yoshioka, K., Fukunaga, R., Sagawa, H., and Matsuura, H. (2018). A simple antegrade perfusion method for isolating viable single cardiomyocytes from neonatal to aged mice. Physiol. Rep. 6, e13688.

Palmer, J.N., Hartogensis, W.E., Patten, M., Fortuin, F.D., and Long, C.S. (1995). Interleukin-1 beta induces cardiac myocyte growth but inhibits cardiac fibroblast proliferation in culture. J. Clin. Invest. 95, 2555–2564.

Pianca, N., Di Bona, A., Lazzeri, E., Costantini, I., Franzoso, M., Prando, V., Armani, A., Rizzo, S., Fedrigo, M., Angelini, A., et al. (2019). Cardiac sympathetic innervation network shapes the myocardium by locally controlling cardiomyocyte size through the cellular proteolytic machinery. J. Physiol. 597.

Pianca, N., Sacchi, F., Umansky, K.B., Chirivì, M., Iommarini, L., Da Pra, S., Papa, V., Bongiovanni, C., Miano, C., Pontis, F., et al. (2022). Glucocorticoid receptor antagonization propels endogenous cardiomyocyte proliferation and cardiac regeneration. Nat. Cardiovasc. Res. 1, 617–633.

Polizzotti, B.D., Ganapathy, B., Walsh, S., Choudhury, S., Ammanamanchi, N., Bennett, D.G., Dos Remedios, C.G., Haubner, B.J., Penninger, J.M., Kuhn, B., et al. (2015). Neuregulin stimulation of cardiomyocyte regeneration in mice and human myocardium reveals a therapeutic window. Sci Transl Med 7, 281ra45.

Porrello, E.R., Mahmoud, A.I., Simpson, E., Hill, J.A., Richardson, J.A., Olson, E.N., and Sadek, H.A. (2011). Transient regenerative potential of the neonatal mouse heart. Science 331, 1078–1080.

Poss, K.D., Wilson, L.G., and Keating, M.T. (2002). Heart regeneration in zebrafish. Science 298, 2188–2190.

Quaife-Ryan, G.A., Sim, C.B., Ziemann, M., Kaspi, A., Rafehi, H., Ramialison, M., El-Osta, A., Hudson, J.E., and Porrello, E.R. (2017). Multi-Cellular Transcriptional Analysis of Mammalian Heart Regeneration. Circulation.

Sadek, H., and Olson, E.N. (2020). Toward the Goal of Human Heart Regeneration. Cell Stem Cell 26, 7–16.

Salido-Medina, A.B., Gil, A., Expósito, V., Martínez, F., Redondo, J.M., Hurlé, M.A., Nistal, J.F., and García, R. (2022). BMP7-based peptide agonists of BMPR1A protect the left ventricle against pathological remodeling induced by pressure overload. Biomed. Pharmacother. 149, 112910.

Sampaio-Pinto, V., Rodrigues, S.C., Laundos, T.L., Silva, E.D., Vasques-Nóvoa, F., Silva, A.C., Cerqueira, R.J., Resende, T.P., Pianca, N., Leite-Moreira, A., et al. (2018). Neonatal Apex Resection Triggers Cardiomyocyte Proliferation, Neovascularization and Functional Recovery Despite Local Fibrosis. Stem Cell Reports 10, 860–874.

Schmid, B., Fürthauer, M., Connors, S.A., Trout, J., Thisse, B., Thisse, C., and Mullins, M.C. (2000). Equivalent genetic roles for bmp7/snailhouse and bmp2b/swirl in dorsoventral pattern formation. Development 127, 957–967.

Schnabel, K., Wu, C.C., Kurth, T., and Weidinger, G. (2011). Regeneration of cryoinjury induced necrotic heart lesions in zebrafish is associated with epicardial activation and cardiomyocyte proliferation. PLoS One 6, e18503.

Senyo, S.E., Steinhauser, M.L., Pizzimenti, C.L., Yang, V.K., Cai, L., Wang, M., Wu, T.-D.D., Guerquin-Kern, J.-L.L., Lechene, C.P., and Lee, R.T. (2013). Mammalian heart renewal by pre-existing cardiomyocytes. Nature 493, 433–436.

Shawi, M., and Serluca, F.C. (2008). Identification of a BMP7 homolog in zebrafish expressed in developing organ systems. Gene Expr. Patterns 8, 369–375.

Shen, H., Gan, P., Wang, K., Darehzereshki, A., Wang, K., Kumar, S.R., Lien, C.-L., Patterson, M., Tao, G., and Sucov, H.M. (2020). Mononuclear diploid cardiomyocytes support neonatal mouse heart regeneration in response to paracrine IGF2 signaling. Elife 9.

Soonpaa, M.H., and Field, L.J. (1998). Survey of studies examining mammalian cardiomyocyte DNA synthesis. Circ Res 83, 15–26.

Soonpaa, M.H., Kim, K.K., Pajak, L., Franklin, M., and Field, L.J. (1996). Cardiomyocyte DNA synthesis and binucleation during murine development. Am. J. Physiol. 271, H2183–9.

Sun, L., Yu, J., Qi, S., Hao, Y., Liu, Y., and Li, Z. (2014). Bone Morphogenetic Protein-10 Induces Cardiomyocyte Proliferation and Improves Cardiac Function after Myocardial Infarction. J. Cell. Biochem.

Sun, Y., Fu, J., Xue, X., Yang, H., and Wu, L. (2018). BMP7 regulates lung fibroblast proliferation in newborn rats with bronchopulmonary dysplasia. Mol. Med. Rep. 17, 6277–6284.

Talman, V., Teppo, J., Pöhö, P., Movahedi, P., Vaikkinen, A., Karhu, S.T., Trošt, K., Suvitaival, T., Heikkonen, J., Pahikkala, T., et al. (2018). Molecular Atlas of Postnatal Mouse Heart Development. J. Am. Heart Assoc. 7, e010378.

Tan, C.Y., Wong, J.X., Chan, P.S., Tan, H., Liao, D., Chen, W., Tan, L.W., Ackers-Johnson, M., Wakimoto, H., Seidman, J.G., et al. (2019). Yin Yang 1 Suppresses Dilated Cardiomyopathy and Cardiac Fibrosis Through Regulation of Bmp7 and Ctgf. Circ. Res. 125, 834–846.

Tang, P., Ma, S., Dong, M., Wang, J., Chai, S., Liu, T., and Li, J. (2018). Effect of interleukin-6 on myocardial regeneration in mice after cardiac injury. Biomed. Pharmacother. 106, 303–308.

Tao, Z., Chen, B., Tan, X., Zhao, Y., Wang, L., Zhu, T., Cao, K., Yang, Z., Kan, Y.W., and Su, H. (2011). Coexpression of VEGF and angiopoietin-1 promotes angiogenesis and cardiomyocyte proliferation reduces apoptosis in porcine myocardial infarction (MI) heart. Proc. Natl. Acad. Sci. U. S. A. 108, 2064–2069.

Tate, M., Perera, N., Prakoso, D., Willis, A.M., Deo, M., Oseghale, O., Qian, H., Donner, D.G., Kiriazis, H., De Blasio, M.J., et al. (2021). Bone Morphogenetic Protein 7 Gene Delivery Improves Cardiac Structure and Function in a Murine Model of Diabetic Cardiomyopathy. Front. Pharmacol. 12, 719290.

Tzahor, E., and Poss, K.D. (2017). Cardiac regeneration strategies: Staying young at heart. Science 356, 1035–1039.

Uygur, A., and Lee, R.T. (2016). Mechanisms of Cardiac Regeneration. Dev. Cell 36, 362–374.

Vukicevic, S., Colliva, A., Kufner, V., Martinelli, V., Moimas, S., Vodret, S., Rumenovic, V., Milosevic, M., Brkljacic, B., Delic-Brkljacic, D., et al. (2022). Bone morphogenetic protein 1.3 inhibition decreases scar formation and supports cardiomyocyte survival after myocardial infarction. Nat. Commun. 13, 81.

Wei, K., Serpooshan, V., Hurtado, C., Diez-Cuñado, M., Zhao, M., Maruyama, S., Zhu, W., Fajardo, G., Noseda, M., Nakamura, K., et al. (2015). Epicardial FSTL1 reconstitution regenerates the adult mammalian heart. Nature 525, 479–485.

Weiskirchen, R., Meurer, S.K., Gressner, O.A., Herrmann, J., Borkham-Kamphorst, E., and Gressner, A.M. (2009). BMP-7 as antagonist of organ fibrosis. Front. Biosci. (Landmark Ed. 14, 4992–5012.

Wodsedalek, D.J., Paddock, S.J., Wan, T.C., Auchampach, J.A., Kenarsary, A., Tsaih, S.-W., Flister, M.J., and O’Meara, C.C. (2019). IL-13 promotes in vivo neonatal cardiomyocyte cell cycle activity and heart regeneration. Am. J. Physiol. Heart Circ. Physiol. 316, H24–H34.

Wu, C.-C., Kruse, F., Vasudevarao, M.D., Junker, J.P., Zebrowski, D.C., Fischer, K., Noël, E.S., Grün, D., Berezikov, E., Engel, F.B., et al. (2016). Spatially Resolved Genome-wide Transcriptional Profiling Identifies BMP Signaling as Essential Regulator of Zebrafish Cardiomyocyte Regeneration. Dev. Cell 36, 36–49.

Yamashita, H., Dijke, P. Ten, Huylebroeck, D., Kuber Sampath, T., Andries, M., Smith, J.C., Heldin, C.H., and Miyazono, K. (1995). Osteogenic protein-1 binds to activin type II receptors and induces certain activin-like effects. J. Cell Biol. 130, 217–226.

Ye, L., D’Agostino, G., Loo, S.J., Wang, C.X., Su, L.P., Tan, S.H., Tee, G.Z., Pua, C.J., Pena, E.M., Cheng, R.B., et al. (2018). Early Regenerative Capacity in the Porcine Heart. Circulation 138, 2798–2808.

Zacchigna, S., Martinelli, V., Moimas, S., Colliva, A., Anzini, M., Nordio, A., Costa, A., Rehman, M., Vodret, S., Pierro, C., et al. (2018a). Paracrine effect of regulatory T cells promotes cardiomyocyte proliferation during pregnancy and after myocardial infarction. Nat. Commun. 9, 2432.

Zacchigna, S., Martinelli, V., Moimas, S., Colliva, A., Anzini, M., Nordio, A., Costa, A., Rehman, M., Vodret, S., Pierro, C., et al. (2018b). Paracrine effect of regulatory T cells promotes cardiomyocyte proliferation during pregnancy and after myocardial infarction. Nat. Commun. 1–12.

Zaidi, S.H.E., Huang, Q., Momen, A., Riazi, A., and Husain, M. (2010). Growth differentiation factor 5 regulates cardiac repair after myocardial infarction. J. Am. Coll. Cardiol. 55, 135–143.

Zeisberg, E.M., Tarnavski, O., Zeisberg, M., Dorfman, A.L., McMullen, J.R., Gustafsson, E., Chandraker, A., Yuan, X., Pu, W.T., Roberts, A.B., et al. (2007). Endothelial-to-mesenchymal transition contributes to cardiac fibrosis. Nat. Med. 13, 952–961.

Zhang, Y.E. (2017). Non-Smad signaling pathways of the TGF-β family. Cold Spring Harb. Perspect. Biol.

Zhu, W., Zhang, E., Zhao, M., Chong, Z., Fan, C., Tang, Y., Hunter, J.D., Borovjagin, A. V., Walcott, G.P., Chen, J.Y., et al. (2018). Regenerative Potential of Neonatal Porcine Hearts. Circulation 138, 2809–2816.

