## Supplementary Figures and Tables for "BMP7 promotes cardiomyocyte regeneration"

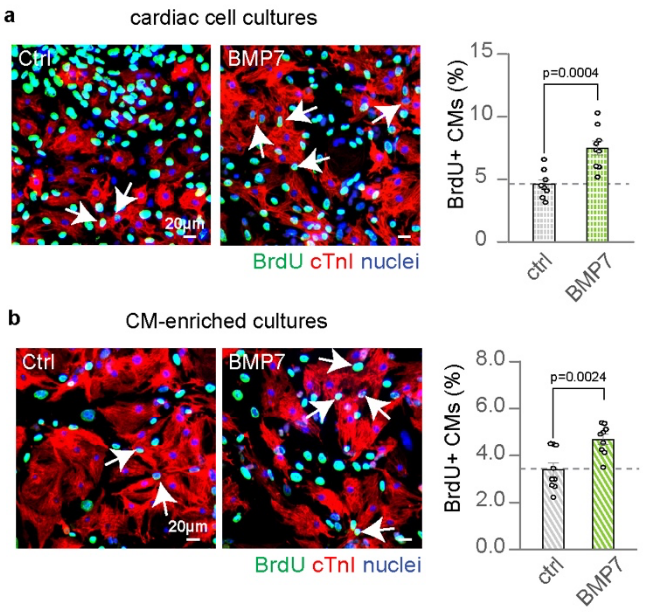


π

**Supplementary Fig. 1. Evaluation of the direct effect of BMP7 on cardiomyocyte proliferation. (a-b)** Immunofluorescence analysis of cardiomyocyte DNA synthesis (BrdU assay) in **(a)** cardiac or **(b)** cardiomyocyte-enriched cell cultures obtained by immunomagnetic separation, isolated from postnatal day 1 (P1) mice, following stimulation with BMP7 at 10 ng/mL for 48 hours (n = 8987 cardiomyocytes pooled from the analysis of 18 samples for cardiomyocyte-enriched cultures and n = 5257 cardiomyocytes pooled from the analysis of 18 samples for cardiac cell cultures); representative pictures in **a-b** are provided; arrows point at proliferating cardiomyocytes; scale bars, 20 µm. The values are presented as mean (error bars show s.e.m.); statistical significance was determined using two-sided Student's t-test in **a** and **b**.

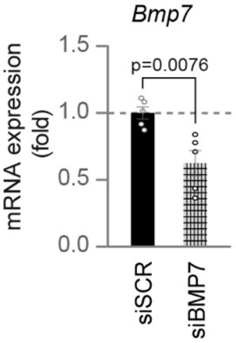


**Supplementary Fig. 2. Analysis of BMP7 knockdown efficacy in cardiomyocyte cultures.** mRNA expression levels of *Bmp7* on postnatal day 1 (P1) cardiac cells, following siRNA delivery for 48 hours (n = 10). The values are presented as mean (error bars show s.e.m.); statistical significance was determined using two-sided Student's t-test.

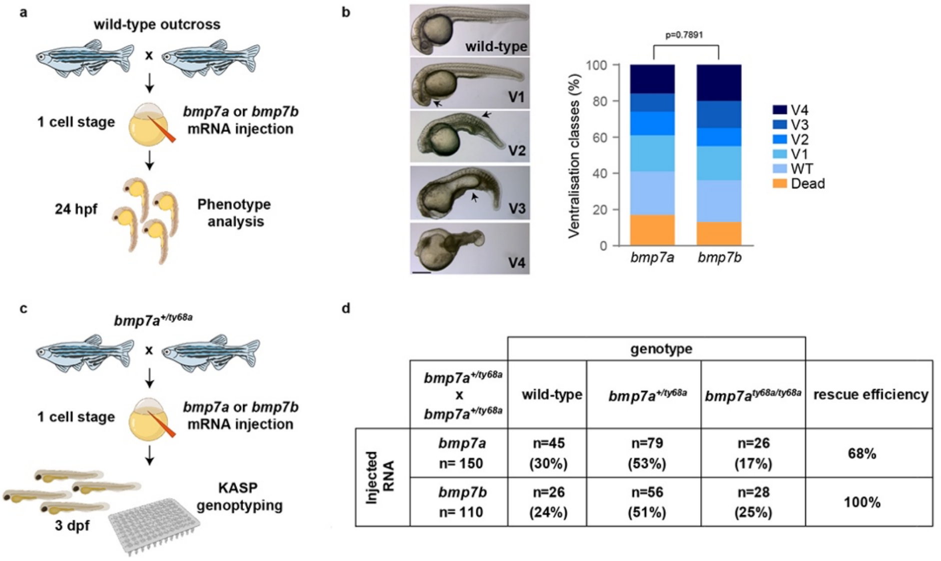


| $bmp7a^{+/ty68a}$ x $bmp7a^{+/ty68a}$ | genotype: wild-type | genotype: $bmp7a^{+/ty68a}$ | genotype: $bmp7a^{ty68a/ty68a}$ | rescue efficiency |
|---|---|---|---|---|
| *bmp7a* n= 150 | n=45 (30%) | n=79 (53%) | n=26 (17%) | 68% |
| *bmp7b* n= 110 | n=26 (24%) | n=56 (51%) | n=28 (25%) | 100% |

**Supplementary Fig. 3. Zebrafish *bmp7a* and *bmp7b* have similar biological activity when overexpressed in embryos.** (**a**) Experimental set-up for assessing phenotypes induced by *bmp7a* and *bmp7b* overexpression during early embryonic development; (**b**) Phenotypes of embryos derived from wild-type outcrosses and injected with 300 pg *bmp7a* or *bmp7b* mRNA. Binning of observed phenotypes into ventralisation classes V1-V4 according to Kishimoto et al., 1997 (for *bmp7a* n = 145, for *bmp7b* n = 150). Representative pictures are provided; scale bars, 100 μm; the graph displays the distribution of embryos in %; (**c**) Experimental set-up for comparing the ability of *bmp7a* and *bmp7b* to rescue the lethal dorsalisation phenotypes of *bmp7a* (*snailhouse*) mutant embryos; (**d**) Genotypes of embryos derived from incrosses of $bmp7a^{ty68a}$ heterozygous parents and injected with *bmp7a* or *bmp7b* mRNA. The expected genotype distribution is 1/3 wild type and 2/3 heterozygous in uninjected clutches, since homozygous mutant embryos die before 3 dpf. In case of complete rescue, the expected genotype distribution is 25% wild-type, 50% heterozygous and 25% homozygous mutant. The rescue efficiency was calculated by dividing the % mutant genotype by the % expected mutant genotype after complete rescue (n = number of genotyped embryos); Statistical significance in **b** was determined using Chi-square test. Some elements in **a** and **c** were created with biorender.com.

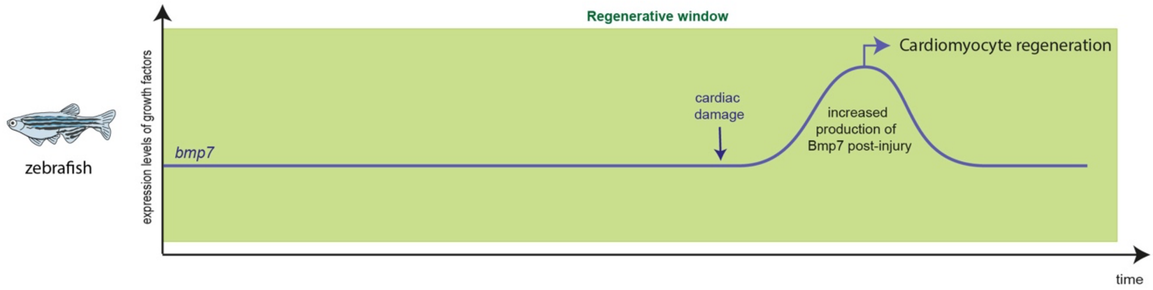


**Supplementary Fig. 4. Model of the role of Bmp7 in cardiomyocyte proliferation and heart regeneration in non-mammalian vertebrates (zebrafish model).** In zebrafish, *bmp7* expression is induced after cardiac injury, in turn triggering cardiomyocyte proliferation and heart regeneration.

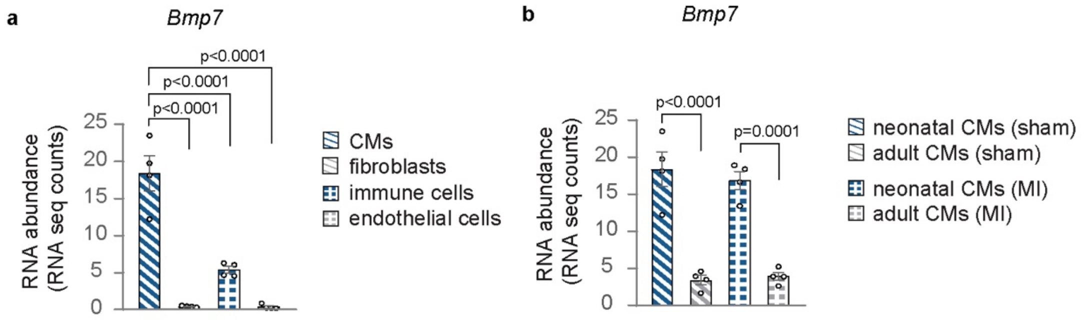


**Supplementary Fig. 5. Expression levels of *Bmp7* in different cardiac cell populations isolated from non-infarcted and infarcted neonatal and adult mice. (a)** Analysis of RNA-sequencing data (Quaife-Ryan et al., 2017) for *Bmp7* expression levels in different cardiac populations (cardiomyocytes, fibroblasts, immune cells and endothelial cells) isolated from neonatal non-infarcted (sham surgery) mice (n = 4 mice); **(b)** Analysis of RNA-sequencing data (Quaife-Ryan et al., 2017) for *Bmp7* expression levels in cardiomyocytes isolated from non-infarcted (sham surgery) and infarcted mice at the neonatal and adult stage (n= 4 mice per condition). The values are presented as mean (error bars show s.e.m.), statistical significance was determined using one way ANOVA followed by Tukey's test.

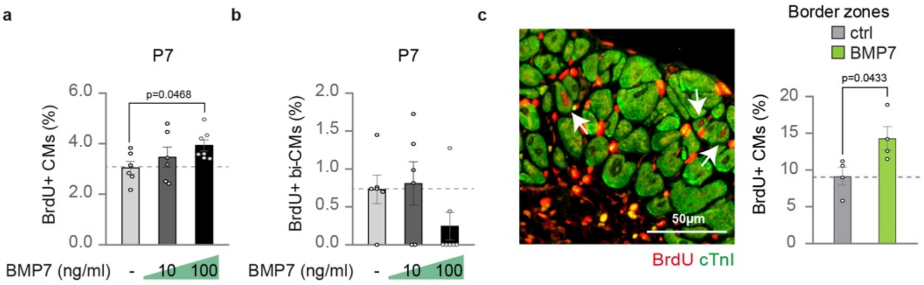


**Supplementary Fig. 6. Mitogenic activity of BMP7 on cardiomyocytes at juvenile stage. (a-b)** Quantification of DNA synthesis (BrdU assay) on postnatal day 7 (P7) cardiomyocytes (total cardiomyocytes in **a** and binucleated cardiomyocytes in **b**) following stimulation with BMP7 at 10 and 100 ng/mL for approximatively 48 hours (n = 2843 cardiomyocytes pooled from the analysis of 19 samples); (**c**) *In vivo* evaluation of adult cardiomyocyte progression to S-phase by BrdU assay and Troponin I (cTnI) immunofluorescence analysis in the border zones of heart sections 14 days post myocardial infarction, following daily injection of BMP7 or water as control (n = 8 mice; a total of 10749 cardiomyocytes have been analysed). BrdU-positive cardiomyocytes in tissue sections were counted manually within border zones; every dot represents a different heart (biological replicate), which in turn has been calculated as average of the analysis of 1 to 2 sections. A representative picture is provided; arrows point at proliferating cardiomyocytes; scale bars 50 µm. The values are presented as mean (error bars show s.e.m.); statistical significance was determined using one way ANOVA followed by Sidak's test by comparing pairs of treatments in **a-b** (control vs selected BMP7 dose) and using two-sided Student's t-test in **c**.

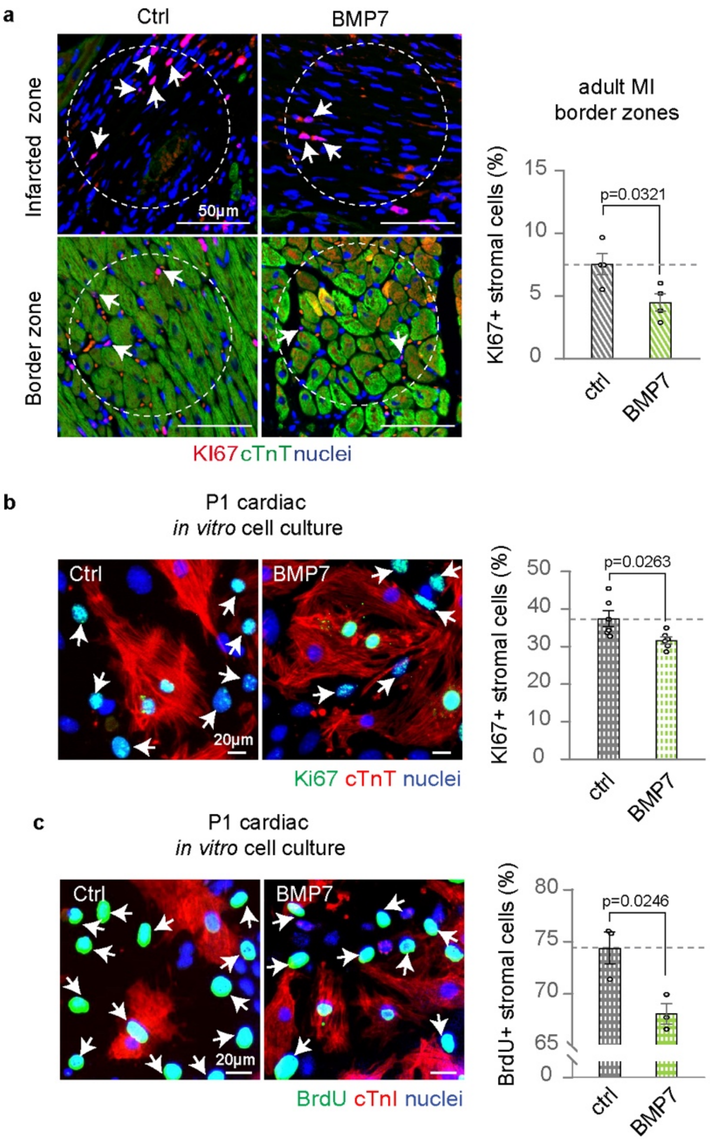


**Supplementary Fig. 7. BMP7 reduces the proliferation of cardiac stromal cells *in vitro* and *in vivo*.** (**a**) Immunofluorescence analysis of cell-cycle activity (KI67) *in vivo* on border zones of heart sections 14 days post myocardial infarction, following daily injections of BMP7 or water as control (n = 8 mice each, with a total of 13683 stromal cells analysed); KI67-positive stromal cells were counted manually within border zones from 12 randomly selected circular areas; each dot represents a different heart (biological replicate), which is the average of the analysis of 2 to 3 sections. Representative pictures are provided; random circular areas are indicated; arrows point at proliferating stromal cells; scale bars, 50 µm; (**b**) Immunofluorescence analysis of cell-cycle re-entry (KI67) *in vitro* on neonatal (postnatal day 1, P1) stromal cardiac cells following stimulation with BMP7 at 10 ng/ml for 48 hours (n = 7914 stromal cells pooled from the analysis of 12 samples); **(c)** Immunofluorescence analysis of cell-cycle progression (BrdU assay) *in vitro* on neonatal (postnatal day 1, P1) stromal cardiac cells following stimulation with BMP7 at 10 ng/ml for 48 hours (n = 5719 stromal cells pooled from the analysis of 6 samples). Representative pictures for **b**-**c** are provided; arrows point at proliferating cardiac stromal cells; scale bars, 20 µm. The values in **a**, **b** and **c** are presented as mean (error bars show s.e.m.), statistical significance was determined using two-sided Student's t-test.

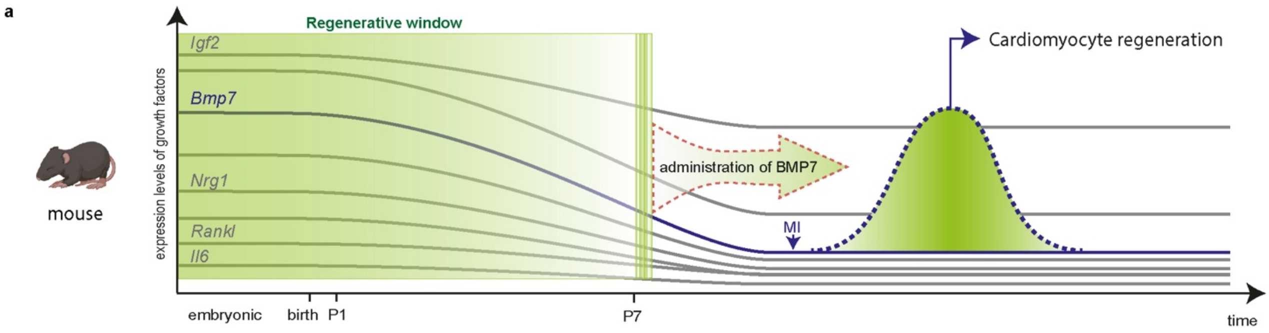


**Supplementary Fig. 8. Model of the role of BMP7 in cardiomyocyte proliferation and heart regeneration in mammals (mouse model).** In mice, the expression levels of several pro-regenerative growth factors (*Il6, Rankl, Nrg1, Bmp7, Igf2*) decline during the first week after birth contributing to cardiomyocyte cell cycle withdrawal. The administration of BMP7 to P7 mice is able to prolong the cardiomyocyte proliferative window, and triggers cardiomyocyte regeneration following myocardial infarction (MI).

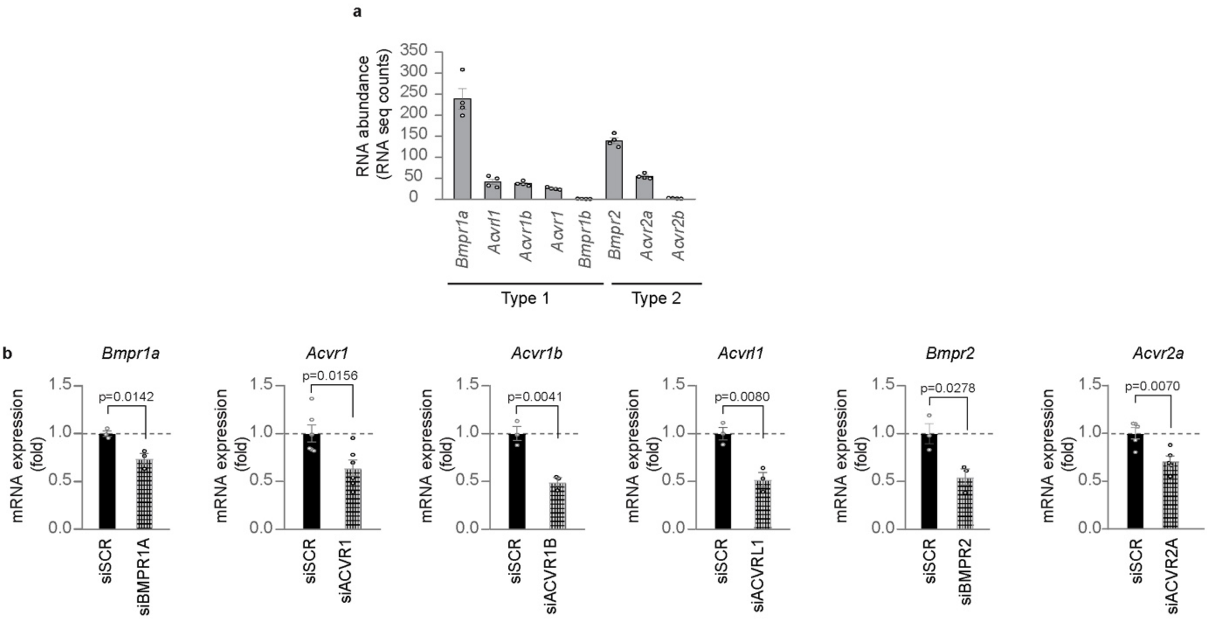


**Supplementary Fig. 9.** ***In vivo*** **evaluation of BMP receptor expression levels in cardiac tissue of neonatal mice and** ***in vitro*** **analysis of the efficacy of BMP receptor gene knockdown in neonatal cardiomyocyte cultures. (a)** Analysis of expression levels of type I *(Bmpr1a*, *Acvrl1*, *Acvr1b*, *Acvr1*, and *Bmpr1b*) and type II (*Bmpr2*, *Acvr2a*, and *Acvr2b*) BMP receptors in neonatal cardiomyocytes of sham surgery (non-infarcted) mice (n = 4 samples) (multicellular RNA sequencing data from (Quaife-Ryan et al., 2017)); **(b)** mRNA expression levels of *Bmpr1a*, *Acvr1*, *Acvr1b*, *Acvrl1*, *Bmpr2* and *Acvr2a* on postnatal day 1 (P1) cardiac cells, following siRNA delivery for 48 hours (n = 6 for *Bmpr1a*, n = 12 for *Acvr1*, n = 6 for *Acvr1b*, n = 6 for *Acvrl1* n = 6 for *Bmpr2*, n = 10 for *Acvr2a)*. The values are presented as mean (error bars show s.e.m.), statistical significance in **b** was determined using two-sided Student's t-test.

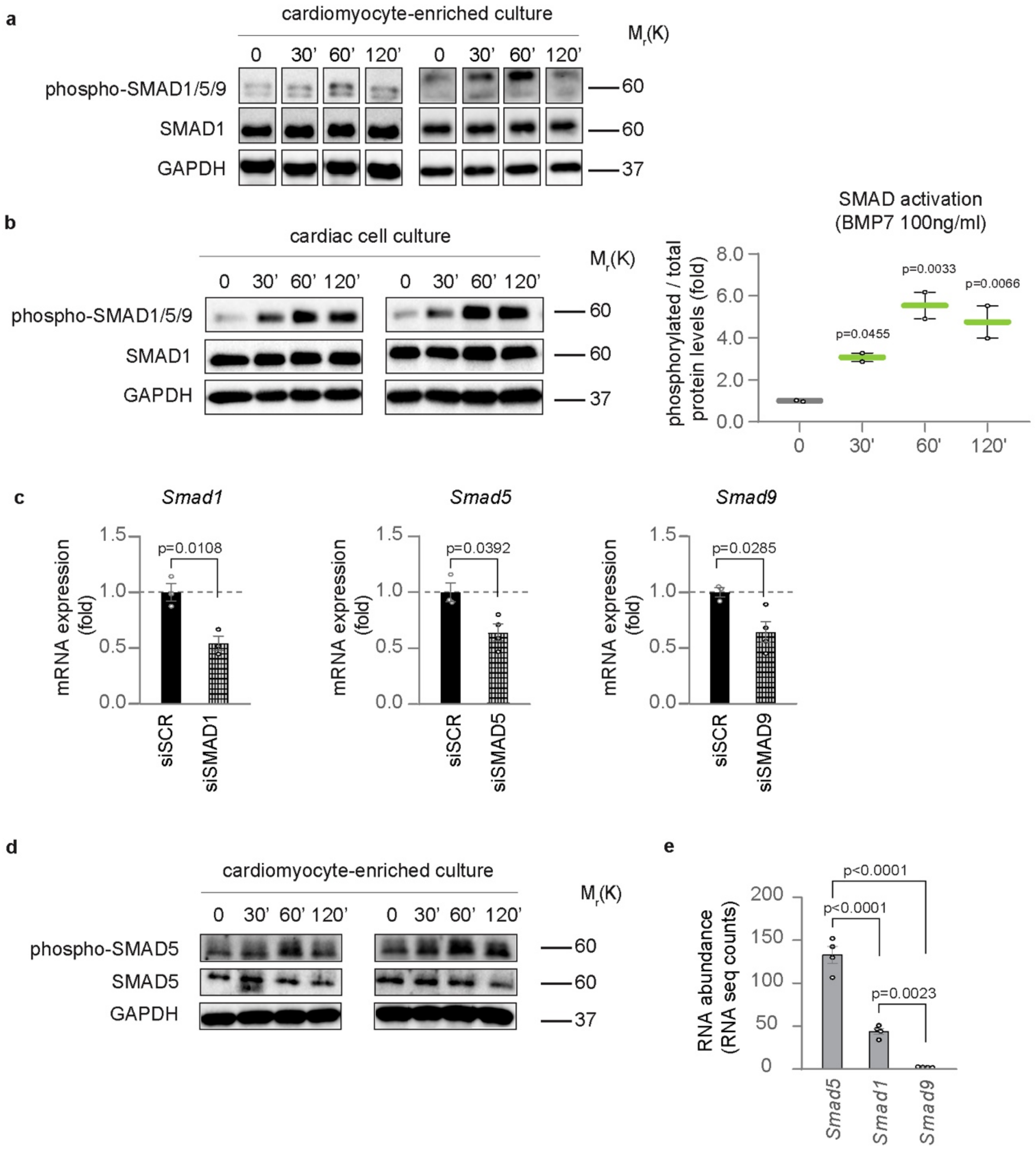


**Supplementary Fig. 10. BMP7 activation of SMAD1/5/9 and analysis of SMAD gene knockdown and expression levels in neonatal cardiomyocyte cultures and in cardiac tissue of neonatal mice, respectively. (a-b)** Western Blot analysis of phospho-SMAD1/5/9, SMAD1 and GAPDH protein levels in **(a)** enriched neonatal (postnatal day 1, P1) cardiomyocytes, separated from stromal cells by immunomagnetic separation (additional 2 sets of replicates included in the graph of Figure 6b), and in **(b)** neonatal (postnatal day 1, P1) cardiac cultures following BMP7 stimulation at 100 ng/ml for 30, 60 and 120 minutes (n = 2 replicates per condition); **(c)** mRNA expression levels of *Smad1*, *Smad5* and *Smad9* on cardiac cells isolated from postnatal day 1 (P1) mice following siRNA delivery for 48 hours (n = 6 for Smad1, n = 7 for Smad5, n = 7 for Smad9); **(d)** Western Blot analysis of phospho-SMAD5, SMAD5 and GAPDH protein levels in enriched neonatal (postnatal day 1, P1)

cardiomyocytes following BMP7 stimulation at 100 ng/ml for 30, 60 and 120 minutes (additional 2 sets of replicates included in the graph of Figure 6d). GAPDH protein levels are provided as second loading control in **a, b** and **d**; (**e**) Analysis of expression levels of SMAD transducers (*Smad1*, *Smad5*, and *Smad9*) in cardiomyocytes isolated from neonatal mice undergoing sham surgery (non-infarcted) (n = 4 samples) (multicellular RNA sequencing data from (Quaife-Ryan et al., 2017)). The values in **b**, **c** and **e** are presented as mean (error bars show s.e.m.), statistical significance was determined using one way ANOVA followed by Sidak's test (comparison between pairs of treatments) for **b,** by Tukey's test for **e** and using two-sided Student's t-test for **c.**

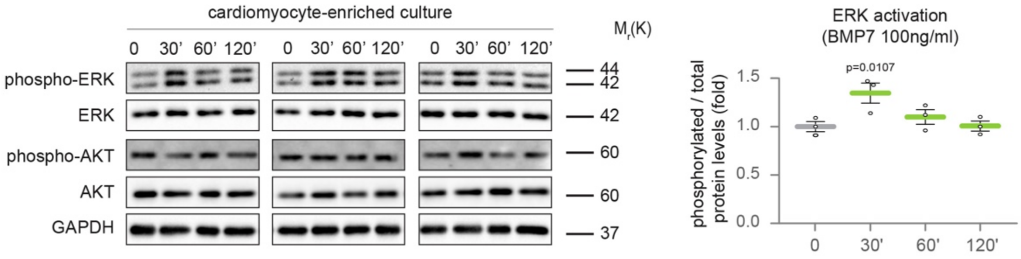


**Supplementary Fig. 11. Analysis of ERK and AKT activation following BMP7 treatment in cardiomyocyte-enriched cultures.** Western Blot analysis of protein levels of phospho-ERK, ERK, phospho-AKT, AKT and GAPDH protein levels in cardiomyocyte-enriched neonatal (postnatal day 1, P1) cultures stimulated with BMP7 at 100 ng/ml for 30, 60 and 120 minutes); The values are presented as mean (error bars show s.e.m.), statistical significance was determined using one way ANOVA followed by Sidak's test (comparison between pairs of treatments).

**Supplementary Table 1.** List of growth factors interrogated for the gene expression profile of hearts in 1-day-old (P1) and 9-10-day-old (P9-P10) mice.

| Growth factor (gene names) | Description |
|---|---|
| *Adipoq* | adiponectin, C1Q and collagen domain containing |
| *Aimp1* | aminoacyl tRNA synthetase complex-interacting multifunctional protein 1 |
| *Areg* | amphiregulin |
| *Artn* | artemin |
| *Bdnf* | brain derived neurotrophic factor |
| *Bmp2* | bone morphogenetic protein 2 |
| *Bmp4* | bone morphogenetic protein 4 |
| *Bmp7* | bone morphogenetic protein 7 |
| *Btc* | betacellulin, epidermal growth factor family member |
| *Ccl11* | chemokine (C-C motif) ligand 11 |
| *Ccl17* | chemokine (C-C motif) ligand 17 |
| *Ccl19* | chemokine (C-C motif) ligand 19 |
| *Ccl2* | chemokine (C-C motif) ligand 2 |
| *Ccl21a* | chemokine (C-C motif) ligand 21A (serine) |
| *Ccl22* | chemokine (C-C motif) ligand 22 |
| *Ccl24* | chemokine (C-C motif) ligand 24 |
| *Ccl25* | chemokine (C-C motif) ligand 25 |
| *Ccl28* | chemokine (C-C motif) ligand 28 |
| *Ccl3* | chemokine (C-C motif) ligand 3 |
| *Ccl4* | chemokine (C-C motif) ligand 4 |
| *Ccl5* | chemokine (C-C motif) ligand 5 |
| *Ccl7* | chemokine (C-C motif) ligand 7 |
| *Cd40* | CD40 antigen |
| *Cntf* | ciliary neurotrophic factor |
| *Csf2* | colony stimulating factor 2 (granulocyte-macrophage) |
| *Csf3* | colony stimulating factor 3 (granulocyte) |
| *Ctgf* | connective tissue growth factor |
| *Ctla4* | cytotoxic T-lymphocyte-associated protein 4 |
| *Cxcl1* | chemokine (C-X-C motif) ligand 1 |
| *Cxcl10* | chemokine (C-X-C motif) ligand 10 |
| *Cxcl11* | chemokine (C-X-C motif) ligand 11 |
| *Cxcl12* | chemokine (C-X-C motif) ligand 12 |
| *Cxcl13* | chemokine (C-X-C motif) ligand 13 |
| *Cxcl14* | chemokine (C-X-C motif) ligand 14 |
| *Cxcl17* | chemokine (C-X-C motif) ligand 17 |
| *Cxcl2* | chemokine (C-X-C motif) ligand 2 |
| *Cxcl5* | chemokine (C-X-C motif) ligand 5 |

| | |
|---|---|
| *Egf* | epidermal growth factor |
| *Epgn* | epithelial mitogen |
| *Ereg* | epiregulin |
| *Fas* | Fas (TNF receptor superfamily member 6) |
| *Fgf1* | fibroblast growth factor 1 |
| *Fgf10* | fibroblast growth factor 10 |
| *Fgf2* | fibroblast growth factor 2 |
| *Fgf23* | fibroblast growth factor 23 |
| *Fgf7* | fibroblast growth factor 7 |
| *Fgf8* | fibroblast growth factor 8 |
| *Flt3l* | FMS-like tyrosine kinase 3 ligand |
| *Fst* | follistatin |
| *Gdf5* | growth differentiation factor 5 |
| *Gdnf* | glial cell line derived neurotrophic factor |
| *Gh* | growth hormone |
| *Grn* | granulin |
| *Hgf* | hepatocyte growth factor |
| *Ifng* | interferon gamma |
| *Igf1* | insulin-like growth factor 1 |
| *Igf2* | insulin-like growth factor 2 |
| *Igfbp1* | insulin-like growth factor binding protein 1 |
| *Igfbp3* | insulin-like growth factor binding protein 3 |
| *Igfbp4* | insulin-like growth factor binding protein 4 |
| *Il1a* | interleukin 1 alpha |
| *Il1b* | interleukin 1 beta |
| *Il1f9* | interleukin 1 family, member 9 |
| *Il1rn* | interleukin 1 receptor antagonist |
| *Il2* | interleukin 2 |
| *Il23a* | interleukin 23, alpha subunit p19 |
| *Il27* | interleukin 27 |
| *Il4* | interleukin 4 |
| *Il5* | interleukin 5 |
| *Il6* | interleukin 6 |
| *Il7* | interleukin 7 |
| *Il10* | interleukin 10 |
| *Il11* | interleukin 11 |
| *Il12b* | interleukin 12b |
| *Il15* | interleukin 15 |
| *Il16* | interleukin 16 |
| *Il17b* | interleukin 17B |
| *Il17f* | interleukin 17F |
| *Il19* | interleukin 19 |

| | |
|---|---|
| *Inhba* | inhibin beta-A |
| *Inhbb* | inhibin beta-B |
| *Kitl* | kit ligand |
| *Lgals1* | lectin, galactose binding, soluble 1 |
| *Lgals3* | lectin, galactose binding, soluble 3 |
| *Lgals7* | lectin, galactose binding, soluble 7 |
| *Lif* | leukemia inhibitory factor |
| *Lta* | lymphotoxin A |
| *Mif* | macrophage migration inhibitory factor (glycosylation-inhibiting factor) |
| *Mstn* | myostatin |
| *Ngf* | nerve growth factor |
| *Nog* | noggin |
| *Nppb* | natriuretic peptide type B |
| *Nrg1* | neuregulin 1 |
| *Ntf3* | neurotrophin 3 |
| *Ntf5* | neurotrophin 5 |
| *Osm* | oncostatin M |
| *Pdgfa* | platelet derived growth factor, alpha |
| *Pdgfb* | platelet derived growth factor, B polypeptide |
| *Pf4* | platelet factor 4 |
| *Pgf* | placental growth factor |
| *Ppbp* | pro-platelet basic protein |
| *Prok1* | prokineticin 1 |
| *Tff2* | trefoil factor 2 (spasmolytic protein 1) |
| *Tgfb1* | transforming growth factor, beta 1 |
| *Tgfb2* | transforming growth factor, beta 2 |
| *Tgfb3* | transforming growth factor, beta 3 |
| *Thpo* | thrombopoietin |
| *Tnf* | tumor necrosis factor |
| *Tnfsf10* | tumor necrosis factor (ligand) superfamily, member 10 |
| *Tnfsf11* | tumor necrosis factor (ligand) superfamily, member 11 |
| *Tnfsf13b* | tumor necrosis factor (ligand) superfamily, member 13b |
| *Tnfsf15* | tumor necrosis factor (ligand) superfamily, member 15 |
| *Tnfsf9* | tumor necrosis factor (ligand) superfamily, member 9 |
| *Tslp* | thymic stromal lymphopoietin |
| *Vegfa* | vascular endothelial growth factor A |
| *Vegfb* | vascular endothelial growth factor B |
| *Vegfc* | vascular endothelial growth factor C |

**Supplementary Table 2.** Sequences of the primers used to analyse mRNA levels by real-time (rt)PCR.

| **Gene** | **Forward primer** | **Reverse primer** |
|---|---|---|
| *Bmpr1a* | 5'-GAAGTATGGATGGGTAAATGG | 5'-ATGTCTGCAGCTATAAAACC |
| *Acvr1* | 5'-GTTGCTCTCAGGAAGTTTAAG | 5'-CTAGTAGTTCCGCTAGAGTG |
| *Acvr2a* | 5'-GGTCTCTTGGAATGAACTTTG | 5'-TTACTTTTGATGTCCCTGTG |
| *Bmpr2* | 5'-GATAATGCGGCTATAAGTGAG | 5'-GTCTGTACACCTCATAAACAC |
| *Bmp7* | 5'-GAAGTCCATCTCCGTAGTATC | 5'-CGTACAGCTCATGTTTCTTG |
| *Hprt1* | 5'-TGGCCGGCAGCGTTTCTGAG | 5'-GTCGGCTCGCGGCAAAAAGC |
| *Acvr1b* | 5'-TCAACATGAAGCACTTTGAC | 5'-GTTGATAGTCTTCATGGACTC |
| *Acvr2b* | 5'-ATTACCTCAAGGGGAACATC | 5'-CATTCTTGCTTTTGAAGTCC |
| *Bmpr1b* | 5'-AATATTCTGGGGTTCATTGC | 5'-CCGTTTTCATGATAGTCTGTG |
| *Acvrl1* | 5'-GAGTGGGTACCAAAAGATA | 5'-GTACCATGTCATAGAAAGGTG |
| *Smad1* | 5'-GGTGCTCTATTGTGTACTATG | 5'-TGGACTCCTTTCCCAATATG |
| *Smad5* | 5'-TATCCAGCAGAGATGTTCAC | 5'-CCAGGCAGAATCTACTTTTG |
| *Smad9* | 5'-TATCAACACTCAGACTTCCG | 5'-GTTTACATTTGAGAGAAGCCC |

**Supplementary Table 3.** Cycling program used for Kompetitive allele specific PCR (KASP) genotyping of zebrafish embryos (LGC Biosearch Technology)

| | |
|---|---|
| Hot-start activation | 94°C, 15 min |
| 10 cycles | 94°C, 20 sec<br>61°C, 60 sec |
| 26 cycles | 94°C, 20 sec<br>55°C, 60 sec |

| | |
|---|---|
| Recycling | |
| 3 cycles | 94°C, 20 sec<br>57°C, 60 sec |
